# Global patterns of subgenome evolution in organelle-targeted genes of six allotetraploid angiosperms

**DOI:** 10.1101/2021.07.09.451712

**Authors:** Joel Sharbrough, Justin L. Conover, Matheus Fernandes Gyorfy, Corrinne E. Grover, Emma R. Miller, Jonathan F. Wendel, Daniel B. Sloan

## Abstract

Whole-genome duplications (WGDs), in which the number of nuclear genome copies is elevated as a result of autopolyploidy or allopolyploidy, are a prominent process of diversification in eukaryotes. The genetic and evolutionary forces that WGD imposes upon cytoplasmic genomes are not well understood, despite the central role that cytonuclear interactions play in eukaryotic function and fitness. Cellular respiration and photosynthesis depend upon successful interaction between the 3000+ nuclear-encoded proteins destined for the mitochondria or plastids and the gene products of cytoplasmic genomes in multi-subunit complexes such as OXPHOS, organellar ribosomes, Photosystems I and II, and Rubisco. Allopolyploids are thus faced with the critical task of coordinating interactions between nuclear and cytoplasmic genes that were inherited from different species. Because cytoplasmic genomes share a more recent history of common descent with the maternal nuclear subgenome than the paternal subgenome, evolutionary “mismatches” between the paternal subgenome and the cytoplasmic genomes in allopolyploids might lead to accelerated rates of evolution in the paternal homoeologs of allopolyploids, either through relaxed purifying selection or strong directional selection to rectify these mismatches. We tested this hypothesis in maternal vs. paternal copies of organelle-targeted genes in six allotetraploids: *Brachypodium hybridum*, *Chenopodium quinoa*, *Coffea arabica*, *Gossypium hirsutum*, *Nicotiana tabacum*, and *Triticum dicoccoides*. We report evidence that allopolyploid subgenomes exhibit unequal rates of protein-sequence evolution, but we did not observe global effects of cytonuclear incompatibilities on paternal homoeologs of organelle-targeted genes. Analyses of gene content revealed mixed evidence for whether organelle-targeted genes re-diploidize more rapidly than non-organelle-targeted genes. Together, these global analyses provide insights into the complex evolutionary dynamics of allopolyploids, showing that allopolyploid subgenomes have separate evolutionary trajectories despite sharing the same nucleus, generation time, and ecological context.

AUTHOR SUMMARY

Whole genome duplication, in which the size and content of the nuclear genome is instantly doubled, represents one of the most profound forms of mutational change. The consequences of duplication events are equally monumental, especially considering that almost all eukaryotes have undergone whole genome duplications during their evolutionary history. While myriad genetic, cellular, organismal, and ecological effects of whole genome duplications have been extensively documented, relatively little attention has been paid to the diminutive but essential “other” genomes present inside the cell, those of chloroplasts and mitochondria. In this study, we compared the evolutionary patterns of >340,000 genes from 23 species to test whether whole genome duplications are associated with genetic mismatches between the nuclear, mitochondrial, and chloroplast genomes. We discovered tremendous differences between duplicated copies of nuclear genomes; however, mitochondria-nuclear and chloroplast-nuclear mismatches do not appear to be common following whole genome duplications. Together these genomic data represent the most extensive analysis yet performed on how polyploids maintain the delicate and finely tuned balance between the nuclear, mitochondrial, and chloroplast genomes.

## INTRODUCTION

Whole genome duplication (WGD) events, in which the nuclear genome is doubled via polyploidization, are among the most profound mutational changes observed in nature. The high frequency of WGDs, especially among flowering plants [1–4], makes them a major force in genome evolution. Accordingly, evolutionary biologists have had a great deal of interest in exploring the consequences of and responses to WGD. The ensuing studies have shown that the effects of WGDs are far-ranging, including silencing and loss of duplicated genes [5–11], mobilization of previously dormant transposable elements [12–17], inter-genomic gene conversion and homoeologous chromosome exchanges [18–25], alterations of epigenetic marks [26–33], massive, genome-wide transcriptional rewiring [6,34–41], and a host of other associated physiological, ecological, and life-history changes [42–54]. Whole genome duplications are also expected to produce novel interactions between the nuclear genome and the mitochondrial and plastid genomes [55], but this dimension of allopolyploid evolution has received relatively little attention (but see [56–62]).

Cytonuclear interactions are themselves the result of gene transfers from the cytoplasmic genomes (mitochondrial and plastid) to the nuclear genome or the recruitment of existing nuclear-encoded proteins to function in these organelles [63, 64]. As a result, the vast majority of the ∼2000 proteins that comprise the mitochondrial proteome [65] and ∼3000 proteins that comprise the plastid proteome [66] are nuclear-encoded [67]. Many of these nuclear-encoded proteins directly interact with gene products from the cytoplasmic genomes to form heteromeric complexes (e.g., Rubisco, Photosystems I and II, organellar ribosomes and the enzymes that comprise the mitochondrial electron transport chain). Additionally, the replication, expression, and post-transcriptional modifications of cytoplasmic genomes are dependent on nuclear-encoded proteins [68–71], as are the many biosynthetic and signaling functions of mitochondria and plastids [72–78]. Taken together, the cellular and metabolic functions that result from cytonuclear interactions, especially aerobic respiration and photosynthesis, are critically important to eukaryotic health and fitness [79–83]. Perturbations to one genomic compartment can therefore have dramatic consequences for the other genomic compartments [84–90], so much so that incompatibilities between the nuclear and cytoplasmic genomes may be a potent force in generating and reinforcing species boundaries [91–95].

Allopolyploidization, a WGD event resulting from a genome merger of two differentiated species [96–98], is expected to perturb cytonuclear interactions because the cytoplasmic genomes have a more recent history of shared descent with one nuclear subgenome than the other [55]. Researchers have hypothesized a number of immediate and evolutionary responses that may mitigate any resulting deleterious consequences. First, maternally biased nuclear gene expression in recently formed allopolyploid lineages could alleviate the deleterious consequences of incompatibilities between the paternal nuclear subgenome and the cytoplasmic genomes [56]. Over time, evolutionary rates may vary across nuclear subgenomes, with paternal copies of organelle-targeted genes evolving faster than maternal copies, either as a reflection of relaxed selection [99] or positive selection to rectify mismatches with the cytoplasmic genomes [90]. In the long run, paternal copies of organelle-targeted genes may be altered more frequently than maternal copies as a result of maternally biased gene conversion [57, 62] and homoeologous exchange [25], or completely excised from the genome via pseudogenization and gene loss [58].

Investigations into the predicted outcomes of cytonuclear incompatibilities in allopolyploids have so far had mixed results. Rubisco in particular has been a primary focus, as the nuclear-encoded small subunit *rbcS* appears to have undergone maternally biased gene conversion and maternally biased gene expression in some allopolyploids, such as cotton, tobacco, *Arabidopsis suecica,* peanut, and wheat [56,57,62]. Synthetic and recently formed allopolyploids show more inconsistent support. For example, *Tragopogon miscellus* exhibits maternally biased expression of *rbcS*, while its reciprocally formed congener *Tragopogon mirus* does not [58]. Synthetic allotetraploid rice showed little evidence of maternally biased expression of *rbcS* [59], and synthetic allopolyploid *Cucumis x hytivus* displayed paternally biased expression of *rbcS* [61]. Generalizing rules of cytonuclear biology from these handful of somewhat contradictory studies is made even more difficult because they have all only considered a single cytonuclear complex.

A more extensive survey of 110 nuclear genes encoding subunits involved in plastid protein complexes in allopolyploid *Brassica napus* did not find evidence for maternally biased expression or retention of organelle-targeted genes [60]. What remains to be evaluated is whether there are systematic rules that might explain the discrepancies among these earlier studies and more generally what the principles are that govern cytonuclear evolution in plant allopolyploids. There are as yet no genome-wide investigations of the signatures of cytonuclear incompatibilities in a set of independently formed allopolyploids that differ both in terms of the amount of divergence between diploid progenitors (and therefore the probability of cytonuclear incompatibilities [100]), or time since allopolyploidization (and therefore the probability of an evolutionary response to cytonuclear incompatibilities [101]). The rapidly increasing availability of genome sequences for a number of allopolyploid genomes and their diploid relatives (e.g., *Brassica* [8,19,102,103], cotton [104–107], wheat [108–112], peanut [23, 113], coffee [114–117], tobacco [118, 119], quinoa [21, 120], and *Brachypodium* [121, 122]) makes it possible to better understand the rules of cytonuclear biology in allopolyploid lineages.

Here we evaluate genome-wide patterns of molecular evolution in organelle-targeted gene sets for six separate allotetraploid species: *Brachypodium hybridum, Chenopodium quinoa* (quinoa), *Coffea arabica* (coffee), *Gossypium hirsutum* (cotton), *Nicotiana tabacum* (tobacco), and *Triticum dicoccoides* (wild emmer wheat). We document strong effects of subgenome on overall rates and patterns of evolution but find little evidence for global signatures of cytonuclear incompatibilities across polyploid systems. We also find that organelle-targeted gene content is generally less biased across subgenomes than the rest of the genome. Together, these genome-wide analyses of six independently formed allotetraploid species provide insights into the rules of polyploidy, a prominent process in eukaryotic diversification.

## RESULTS

### Orthologous genes in six allopolyploid species and their diploid relatives

To compare rates and patterns of molecular evolution across subgenomes of six allotetraploid angiosperms (Figure 1a), we inferred orthologous gene groups from the two polyploid subgenomes, the closest available diploid species for each subgenome, and an outgroup (Figure 2) using a combination of phylogenetic and syntenic methods. The resulting orthologous gene groups are summarized in Table 1, and additional details regarding their inference are provided in the Materials & Methods section as well as in Figure S1.

**Figure 1.**
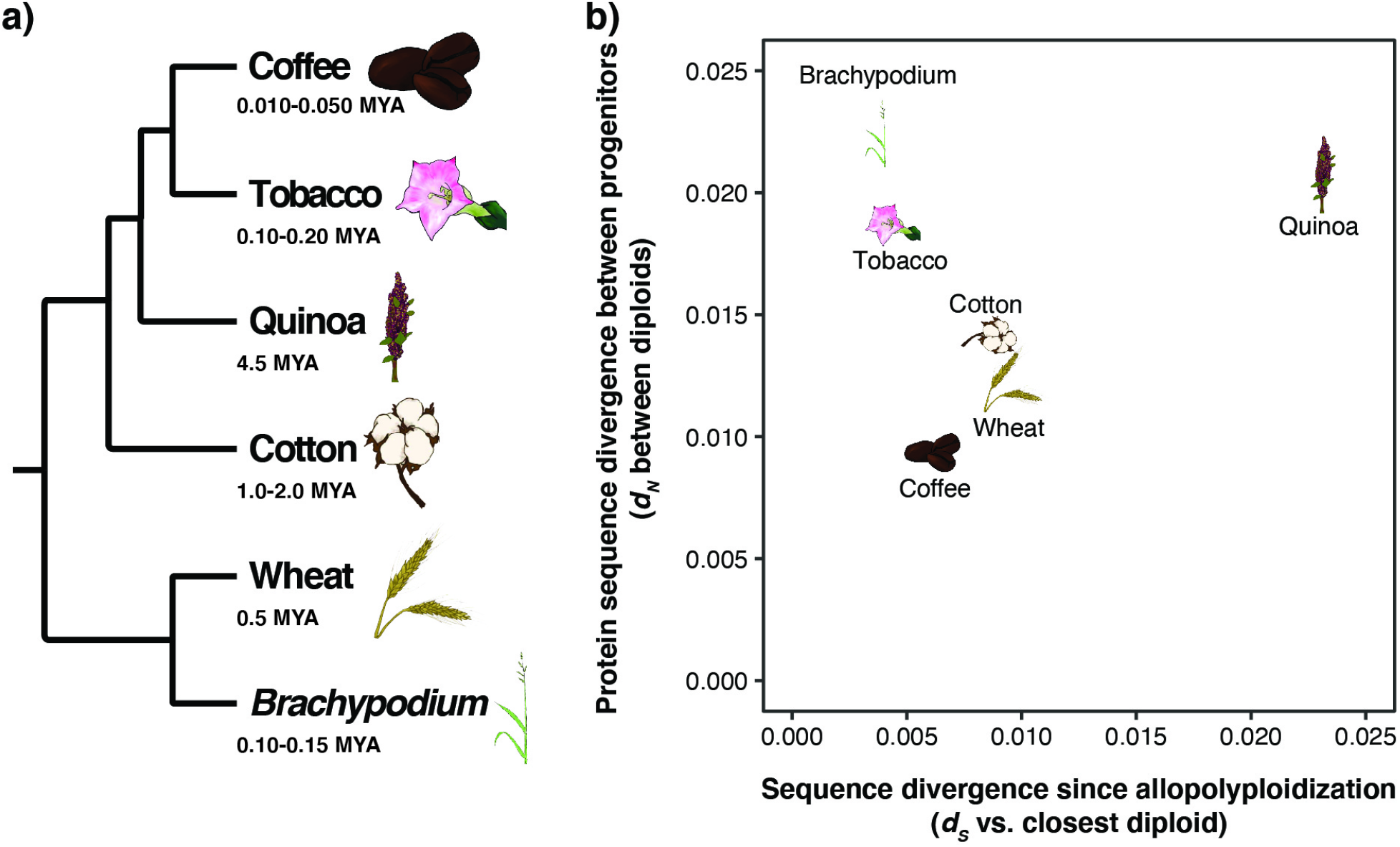
Evolutionary relationships and origins of six allotetraploid angiosperms. a) Cladogram depicting evolutionary relationships among six independently derived allotetraploid angiosperms. b) The scatter plot depicts the synonymous substitutions per synonymous site (*d_S_*) between the polyploid subgenome-diploid pair with the lowest amount of divergence on the x-axis as a proxy for the amount of time since allopolyploidization. Amino acid sequence divergence between subgenomes, measured as nonsynonymous substitutions per nonsynonymous site (*d_N_*) between the two diploid relatives, is shown on the y-axis. Higher levels of amino acid sequence divergence between subgenomes increase the probability of a genetic incompatibility in the polyploid, whereas longer periods of time since allopolyploidization increases the probability that evolutionary responses to incompatibilities are detectable in the polyploid.

**Figure 2.**
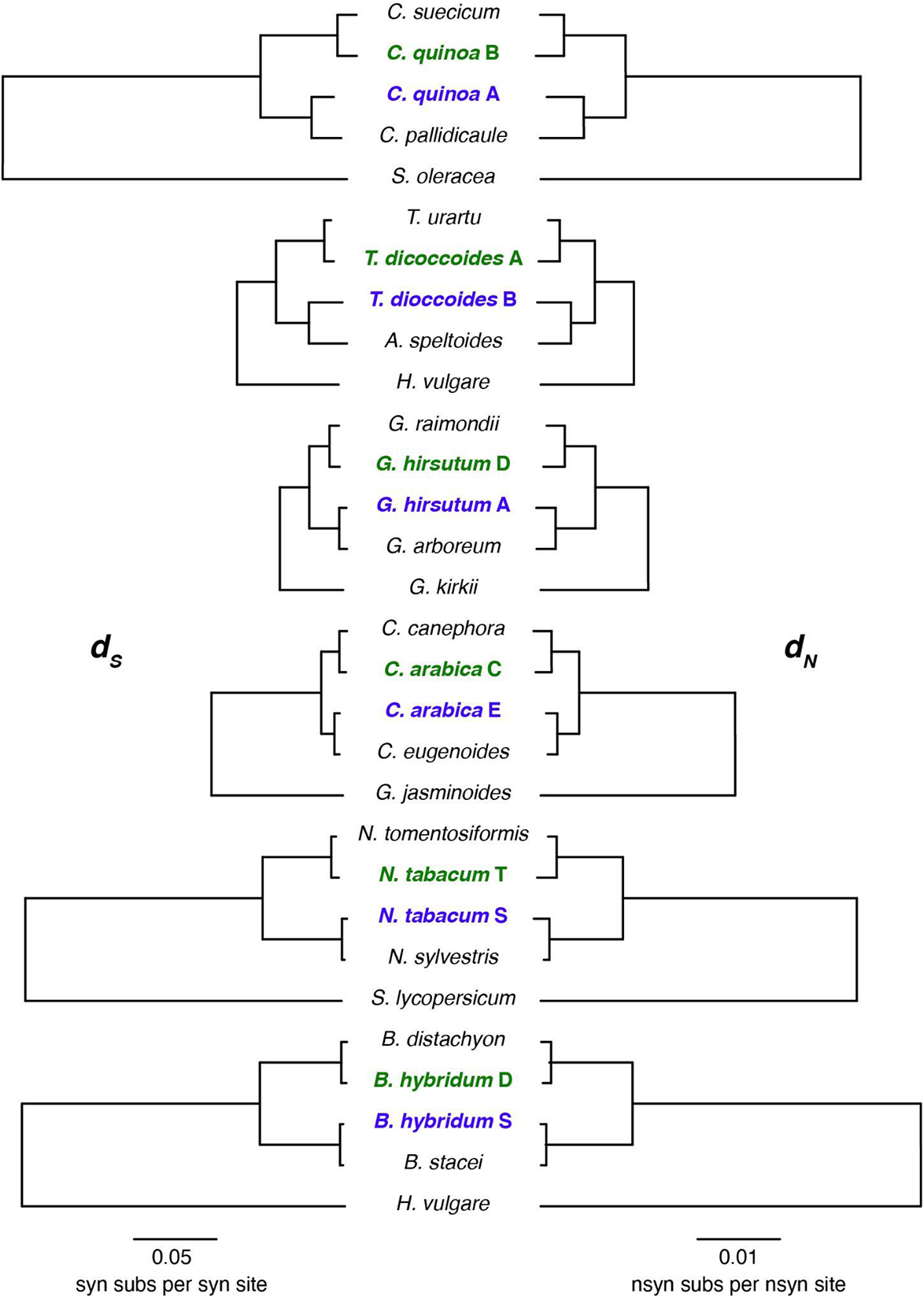
Synonymous and nonsynonymous rates of evolution in genomes (and subgenomes) of focal allopolyploid systems. Substitution rates per site for synonymous (*d_S_* – left) and nonsynonymous (*d_N_* – right) sites from concatenated analyses of non-organelle-targeted genes are represented by branch lengths for each genome (and subgenome). Allopolyploid systems are arranged from oldest (top) to youngest (bottom), as described on the x-axis of Figure 1. Paternal subgenomes of allotetraploids are bolded in green, and maternal subgenomes are bolded in purple.

**Table 1.** Orthologous gene groups in six allotetraploid angiosperms.

| Species | Phylogenetic orthologous gene groups | Syntenic orthologous gene groups | Merged, single-copy quintets<sup>1</sup> (Union/Intersect) | Filtered, merged, single-copy quintets (Union/Intersect) |
| --- | --- | --- | --- | --- |
| Quinoa | 10511 | 17896 | (9088 / 3284) | (8201 / 3121) |
| Wheat | 25454 | 24212 | (10180 / 3603) | (4204 / 1759) |
| Cotton | 29504 | 31841 | (18715 / 10222) | (17133 / 10023) |
| Coffee | 19399 | 20926 | (6672 / 3869) | (4032 / 2379) |
| Tobacco | 24797 | 32088 | (9059 / 167) | (8699 / 163) |
| *Brachypodium* | 24854 | 34440 | (14449 / 8084) | (14000 / 7912) |

<sup>1</sup> – Single-copy quintets include orthologous gene groups with one and only one sequence from an outgroup, two closely related diploids, and two sequences from the allopolyploid.

The goal of our orthology inference methods was to produce orthologous “quintets”, containing one gene sequence each from the outgroup species and the two diploid model species and two gene sequences from the polyploid species, while also requiring that gene trees be consistent with the overall species tree. Both syntenic and phylogenetic methods produced sizable numbers of identical quintets; however, there were many quintets only detectable using one method or the other. Tobacco was especially challenging for syntenic inference, as the highly fragmented assemblies of all three *Nicotiana* reference genomes made identifying syntenic blocks difficult. The largest syntenic block between any two of the genomes in this clade was only 57 genes long (*N. tabacum* and *Solanum lycopersicum*), and no syntenic block including *N. tomentosiformis* or *N. sylvestris* was longer than 22 genes. Quinoa highlighted a different issue that represents a common feature of polyploid genome assemblies in that many genes were located on contigs that are not anchored to chromosomes. Genes present in this fraction of the assembly can only be included in orthologous groups by phylogenetics, and they are often replete with repetitive elements, making it a likely spot for genome misassemblies (and subsequent errors in analyses that depend upon them). Moreover, the quinoa genome contains cases of apparent homoeologous exchange in which genes were located on chromosomes from opposing subgenomes (see also [21]).

Variation in assembly and annotation quality also represented a significant challenge in identifying orthologous genes across genome assemblies produced by different groups with different underlying data. The most extreme example of this issue was the maternal diploid model for polyploid wheat, *Aegilops speltoides*, which was represented only by a transcriptome assembly. Despite these and other hurdles, we were able to identify orthologous gene groups as well as the more strict group of single-copy quintets for each of these polyploid systems, which should present a useful resource for polyploid genomics moving forward. The *Aegilops speltoides* transcriptome, all OrthoFinder results, phylogenetic gene trees with branch lengths, multi-species synteny networks, merged orthologous gene groups, CDS alignments, and analyses of molecular evolution have been made available at https://doi.org/10.6084/m9.figshare.13473207. For the remainder of the manuscript, we report only on the results from the “Union” group of quintets that were identified by either phylogenetic or syntenic inference, but we have performed all the same analyses on the “Intersection” group, comprised only of those quintets that were identified by both methods, and have provided the results from those analyses in Supplementary File 1. Results obtained using the Intersection dataset did not substantively differ from those obtained using the Union dataset.

### Subgenomic distributions of organelle-targeted genes

To evaluate whether cytonuclear interactions affect subgenomic evolution in allopolyploid species, we first partitioned genes by predicted subcellular targeting localization and cytonuclear interaction activity in each allopolyploid system. The Cytonuclear Molecular Interactions Reference for *Arabidopsis* (CyMIRA) database indicates that the *Arabidopsis thaliana* nuclear genome has 1773 genes that encode mitochondria-targeted products and 2931 genes that encode plastid-targeted products [67]. By propagating this classification across the six allotetraploids studied here, we found means of 3880 (SD = 730) genes with mitochondria-targeted products and 4464 (SD = 731) genes with plastid-targeted products (Table 2), which varies ∼60-70% among allotetraploid taxa. At least some of the observed variation among polyploids appears to be due to phylogeny, as the number of mitochondria-targeted genes and plastid-targeted genes varies extensively among diploids (25-30%, Figure S2). Diploid relatives of our focal allotetraploids ranged from 17% fewer (*Chenopodium* diploids) to 108% more (*Nicotiana* diploids) mitochondria-targeted genes and from 37% fewer (*Triticum*, *Chenopodium* diploids) to 33% more (*Nicotiana* diploids) plastid-targeted genes than documented in *Arabidopsis* (Figure S2).

**Table 2.** Functional gene partitioning in six allotetraploid angiosperms.

| Species | Mitochondria-targeted | Mitochondria-targeted Interacting<sup>1</sup> | Mitochondria Enzyme Complexes<sup>2</sup> | Plastid-targeted | Plastid-targeted Interacting<sup>1</sup> | Plastid Enzyme Complexes<sup>2</sup> |
| --- | --- | --- | --- | --- | --- | --- |
| Quinoa | 2830 | 894 | 279 | 3528 | 686 | 215 |
| Wheat | 4077 | 1048 | 378 | 4419 | 693 | 245 |
| Cotton | 4728 | 1232 | 458 | 5670 | 800 | 307 |
| Coffee | 3221 | 921 | 285 | 3889 | 621 | 193 |
| Tobacco | 3851 | 1092 | 402 | 4567 | 740 | 297 |
| *Brachypodium* | 4540 | 981 | 339 | 4684 | 674 | 238 |
| **Mean (SD)** | **3880 (730)** | **1031 (121)** | **358 (68)** | **4464 (731)** | **704 (61)** | **250 (45)** |
| *Arabidopsis thaliana* (diploid) | 1773 | 617 | 180 | 2931 | 375 | 128 |

<sup>1</sup> – Mitochondria- and plastid-targeted interacting genes are a subset of the total number of mitochondria- and plastid-targeted genes
<sup>2</sup> – Mitochondria and plastid enzyme complex genes are a subset of the total number of mitochondria- and plastid-targeted interacting genes

Among the genes with mitochondria-targeted products, CyMIRA lists 617 *A. thaliana* genes that have interactions with mitochondrial genes or gene products and 180 genes with products that are directly involved in enzyme complexes with mitochondrially encoded subunits (i.e., mitoribosome, OXPHOS complexes, TAT complex). We expected to find roughly twice as many genes in each functional category for tetraploids as are present in *Arabidopsis*. In the six focal allotetraploids, we found that functional categories were increased 40-250% (per category/species) relative to *A. thaliana*, with means of 1031 (SD = 121) genes having interactions with mitochondrial genes or gene products and 358 (SD = 68) genes with products that are directly involved in those three mitochondrial enzyme complexes. A similar pattern was observed for genes with plastid-targeted products. Where CyMIRA lists 375 *A. thaliana* genes that have interactions with plastid genes or gene products and 128 genes with products that are directly involved in enzyme complexes with plastid-encoded subunits (i.e., chlororibosome, Photosystems I and II, NDH, ATP synthase, Cytochrome b6f, Rubisco, Clp protease, ACCase), we found means of 704 (SD = 61) and 250 (SD = 45) genes in the allotetraploids for those categories, respectively. Gene numbers for all 55 functional gene categories are described in Table S1, gene IDs for each category and *de novo* targeting predictions are available at https://github.com/jsharbrough/CyMIRA_gene_classification/tree/master/Species_CyMI RA, and the physical distribution of organelle-targeted genes along polyploid chromosomes are shown in Figure S3.

Polyploidization events are expected to perturb cytonuclear interactions in part because cytoplasmic genomes suddenly exist inside a cell in which all of their nuclear-encoded interacting partners have been doubled. One possible evolutionary response to altered cytonuclear stoichiometry in the wake of whole genome duplication is that nuclear-encoded organelle-targeted genes experience selection to rapidly return to a diploid-like state [123, 124]. We tested this hypothesis for both mitochondria- and plastid-targeted nuclear genes in six independently formed allopolyploids using the combined diploid relatives as models for the ancestral allopolyploid state. We found that quinoa (χ^2^ = 54.40, p < 0.0001), wheat (χ^2^ = 660.23, p < 0.0001), tobacco (χ^2^ = 243.85, p < 0.0001), and *Brachypodium* (χ^2^ = 50.15, p < 0.0001) retain a significantly smaller proportion of organelle-targeted genes in duplicate than non-organelle-targeted genes, whereas, cotton (χ^2^ = 134.12, p < 0.0001) and coffee (χ^2^ = 13.40, p = 0.00025) exhibit the opposite pattern by retaining a significantly larger proportion of organelle-targeted genes than non-organelle-targeted genes (Table S2). Notably, the excess retention of organelle-targeted genes in cotton was also evident when we restricted our analysis to only include the subset of genes directly involved in mitochondrial (*χ^2^* = 7.90, *p* = 0.0049) or plastid enzyme complexes (*χ^2^* = 5.58, *p* = 0.018). Although levels of retention within each category varied among species, we did not find a difference in retention levels between mitochondria-targeted versus plastid targeted genes in any of the six species (*p* > 0.05 for all species). Wheat (χ^2^ = 18.35, *p* < 0.0001) and cotton (χ^2^ = 11.05, *p* = 0.00089) both exhibited significantly more pentatricopeptide repeat (PPR) genes (relative to non-organelle-targeted genes) compared to the combined diploids, while the tobacco genome encoded significantly fewer PPR genes than expected (relative to non-organelle-targeted genes) compared to the combined diploids (χ^2^ = 68.09, *p* < 0.0001). Together, these results provide mixed support for rapid loss of organelle-targeted genes compared to the rest of the genome in allopolyploids, but do indicate that similar forces may equally affect mitochondria- and plastid-targeted genes.

A second possible consequence of polyploidy is incompatibility between the paternally derived nuclear subgenome and the maternally derived cytoplasmic genomes, potentially resulting in preferential loss of paternally-derived organelle-targeted genes in hybrid (allo)polyploid species. This effect could exaggerate a general subgenome bias for paternal loss or partially compensate for a general bias towards maternal loss. For five of the allotetraploid genomes, it was possible to assign genes to subgenomes based on their chromosomal position, thereby permitting a relative assessment of parental gene loss; the sole exception, *N. tabacum*, has experienced extensive genomic rearrangement since polyploidization that precludes subgenome assignment based on physical location. In general, we found significant differences in non-organelle-targeted gene abundance across subgenomes for all five allotetraploid species (Table 3), with quinoa, wheat, and coffee exhibiting more paternal homoeolog loss, whereas cotton and *Brachypodium* exhibit a deficit in maternal homoeologs (Figure 3, left panel). Interestingly, however, when considering biases in organelle-targeted genes after correcting for genome-wide levels, these biases flip for quinoa, wheat, and *Brachypodium*. That is, while both quinoa and wheat exhibit biased loss of *paternal* homoeologs for non-organellar targeting genes, those that are targeted to the organelles exhibit biased *maternal* loss (again, relative to background; Figure 3 right panels, Table S3). Similarly, *Brachypodium* organelle-targeted genes exhibit biased paternal loss (relative to background), whereas the genome-wide pattern shows biased maternal loss. These patterns were also found using the diploid relatives to correct for different gene abundances at the time of allopolyploidization (Figure S4). While the maternal homoeolog deficit for organelle-targeted genes found in wheat and quinoa is contrary to predictions based on cytonuclear incompatibilities, we note that this reflects homoeolog retention relative to the genome-wide rate and suggests that these species exhibit a lower degree of subgenomic bias in their organelle-targeted genes than the genome-wide rate.

**Figure 3.**
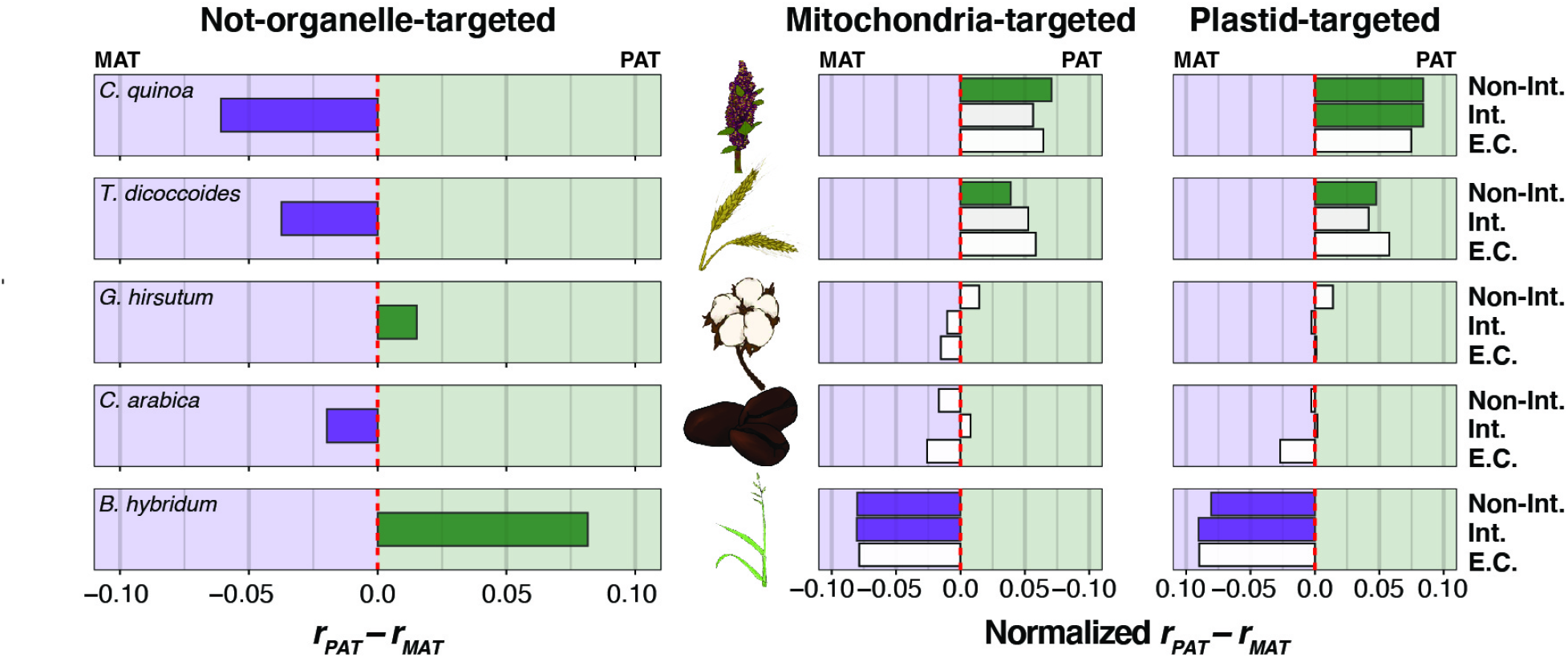
Gene content bias across allotetraploid subgenomes. The proportion of genes present in paternal (*r_PAT_*) vs. maternal (*r_MAT_*) subgenomes is depicted for each of five allotetraploid species arranged vertically from oldest (top) to youngest (bottom). Tobacco was excluded from this analysis because the massive rearrangement it has experienced makes subgenomic identification based on chromosomal position intractable. The left panel includes only non-organelle-targeted genes, the middle panel includes only mitochondria-targeted genes, and the right panel includes only plastid-targeted genes. In the left panel, the red-dashed line represents equal content across subgenomes. In the right two panels, the *r_PAT_* and *r_MAT_* are normalized by the proportions estimated from genes not targeted to the organelles, such that the red-dashed line reflects genome-wide patterns, rather than equality across subgenomes. Proportion deltas that depart significantly from the red line are filled in solid according to the direction of subgenomic bias (i.e., green: *r_PAT_* > *r_MAT_*; purple: *r_PAT_* < *r_MAT_*; no fill: *r_PAT_* ≈ *r_MAT_*). The intimacy of interactions are depicted on the y-axis for each of the right two panels from low- or no-interaction with organelle gene products (top), to interacting genes (middle), to genes involved in mitochondrial or plastid enzyme complexes (bottom).

**Table 3.** Biased gene content of non-organelle-targeted genes across subgenomes of five allotetraploid angiosperms.

| Species | Paternal subgenome | Maternal subgenome | $r_{PAT} - r_{MAT}$ (95% CI) | Binomial test p value |
| --- | --- | --- | --- | --- |
| Quinoa | 9786 | 11053 | -0.061 (-0.074 – -0.047) | < 0.0001 |
| Wheat | 48786 | 52571 | -0.037 (-0.044 – -0.031) | < 0.0001 |
| Cotton | 29762 | 28871 | 0.015 (0.007 – 0.023) | 0.00024 |
| Coffee | 19008 | 19773 | -0.020 (-0.030 – -0.010) | 0.00011 |
| *Brachypodium* | 34860 | 29605 | 0.082 (0.074 – 0.089) | < 0.0001 |

### Evolutionary rate differences across subgenomes and gene functional categories

We used the CyMIRA gene classifications from the maternal diploid models of each allotetraploid to classify single-copy orthologous quintets into functional gene categories, except in the case of wheat. For wheat, the paternal diploid model, *Triticum urartu*, was used because the maternal diploid model (i.e., *Aegilops speltoides*) is only represented by a transcriptome. These functional categories served as the basis for our concatenated and gene-level analyses of evolutionary rate. Summary statistics describing the number of orthologous quintets in each functional category are presented for each allopolyploid system in Table 4 and Figure S5, along with the rates of synonymous (*d*_S_) and nonsynonymous (*d*_N_) evolution in concatenated alignments.

**Table 4.**
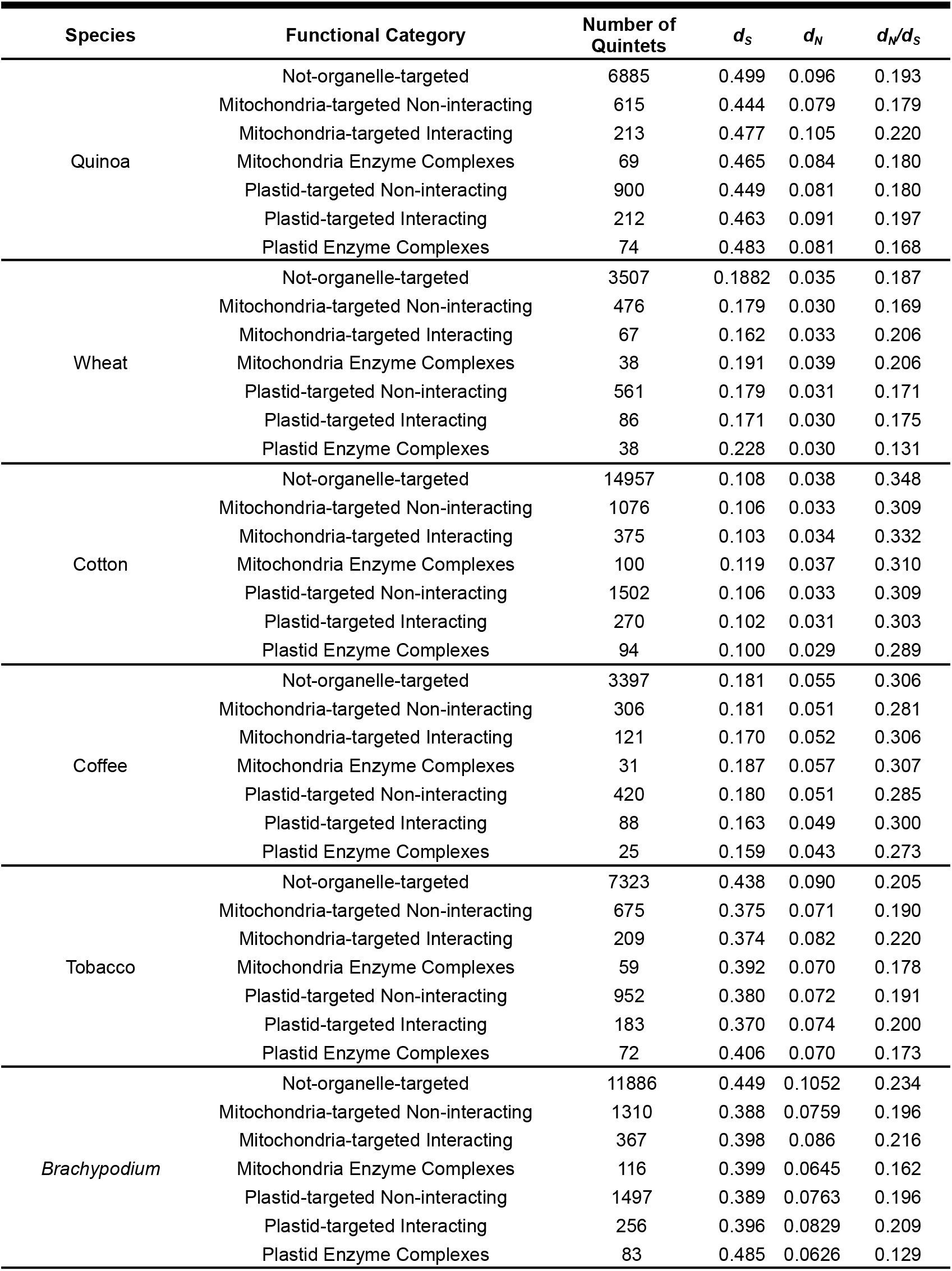
Single-copy orthologous quintets partitioned by functional category in six allotetraploid species.

Rates of protein-sequence evolution vary substantially across CyMIRA functional categories, likely indicative of variation in functional constraint (Figure S5a). In particular, protein sequences of mitochondrial OXPHOS complexes, several of the plastid photosynthesis complexes (but not all, see e.g., the NADH dehydrogenase-like [NDH] complex), as well as the mitochondrial and plastid RNA polymerases appear to evolve especially slowly, indicating that they have experienced relatively stringent purifying selection in these angiosperms. In addition to complex-level effects, we also observed differences in protein-sequence evolution across our focal angiosperm systems, with coffee and cotton genomes exhibiting higher quintet-wide *d_N_/d_S_* values compared to quinoa, wheat, tobacco, and *Brachypodium* (Figure S5b).

Cytonuclear incompatibilities between maternally derived cytoplasmic genomes and the paternal subgenomes of allopolyploids are expected to result in accelerated rates of protein-sequence evolution in the paternal homoeologs of organelle-targeted genes. We tested for signatures of these cytonuclear incompatibilities first by estimating differences in rates of protein-sequence evolution (i.e., *d_N_/d_S_* = *ω*) in concatenated and individual gene alignments of paternal (*ω_PAT_*) vs. maternal (*ω_MAT_*) subgenomes in non-organelle-targeted (NOT) genes to assess whether genome-wide biases exist in our six focal allopolyploids. In concatenated analyses, quinoa, wheat, cotton, and tobacco all showed significant departures (i.e., < 2.5% overlap of bootstrap distributions between *ω_PAT_* and *ω_MAT_*) from equal rates of evolution across subgenomes. In particular, quinoa, cotton, and tobacco exhibited higher *ω* values in maternally derived homoeologs of NOT genes than paternal homoeologs (i.e., *ω_PAT_* : *ω_MAT_* ratio < 1), while coffee and wheat showed the opposite pattern in which paternally derived homoeologs exhibit faster rates of protein-sequence evolution than maternal homoeologs (i.e., *ω_PAT_* : *ω_MAT_* ratio > 1; Figure 4a). We observed similar patterns in gene-level analyses as compared to concatenated analyses in the three older polyploids (Figure 4b): a significantly higher proportion of maternal homoeologs (*p_MAT_*) exhibited faster rates of evolution than paternal homoeologs (*p_PAT_*) in quinoa (binomial test, *p =* 0.0022) and cotton (binomial test, *p* < 0.0001), while *p_PAT_* was significantly greater than *p_MAT_* in wheat (binomial test, *p* < 0.0001). Although *p_MAT_* was greater than *p_PAT_* in the concatenated analysis of tobacco subgenomes, the difference was not significant at the gene level (binomial test, *p =* 0.183). A similar result was obtained in coffee, with the concatenated analysis showing a significant paternal bias, but gene-level patterns did not appear to be paternally biased (binomial test, *p* = 0.375). Bootstrap distributions of *ω_MAT_* in *Brachypodium* estimated from concatenated alignments were higher than bootstrap distributions of *ω_PAT_* but were not significantly different (i.e., > 2.5% overlap), while *p_MAT_* was significantly greater than *p_PAT_* at the individual gene level (binomial test, *p* = 0.00026). The higher *ω* values in the maternal subgenomes of quinoa, cotton, and *Brachypodium* and the higher *ω* values in the paternal subgenome of coffee were primarily driven by differences in *d_N_* as opposed to *d_S_* (Figure 2), indicating that these subgenomes experience different rates of protein-sequence evolution. By contrast, the elevated *ω* values in the maternal subgenome of tobacco and the paternal subgenome of wheat were primarily driven by *d_S_* (Figure 2), potentially indicating that different subgenomes experience different mutation rates or that the diploids used here represent highly asymmetric models of the diploid progenitors. Taken together, these analyses of NOT genes indicate that allopolyploids experience significant biases in rates of evolution across subgenomes present inside the same cell.

**Figure 4.**
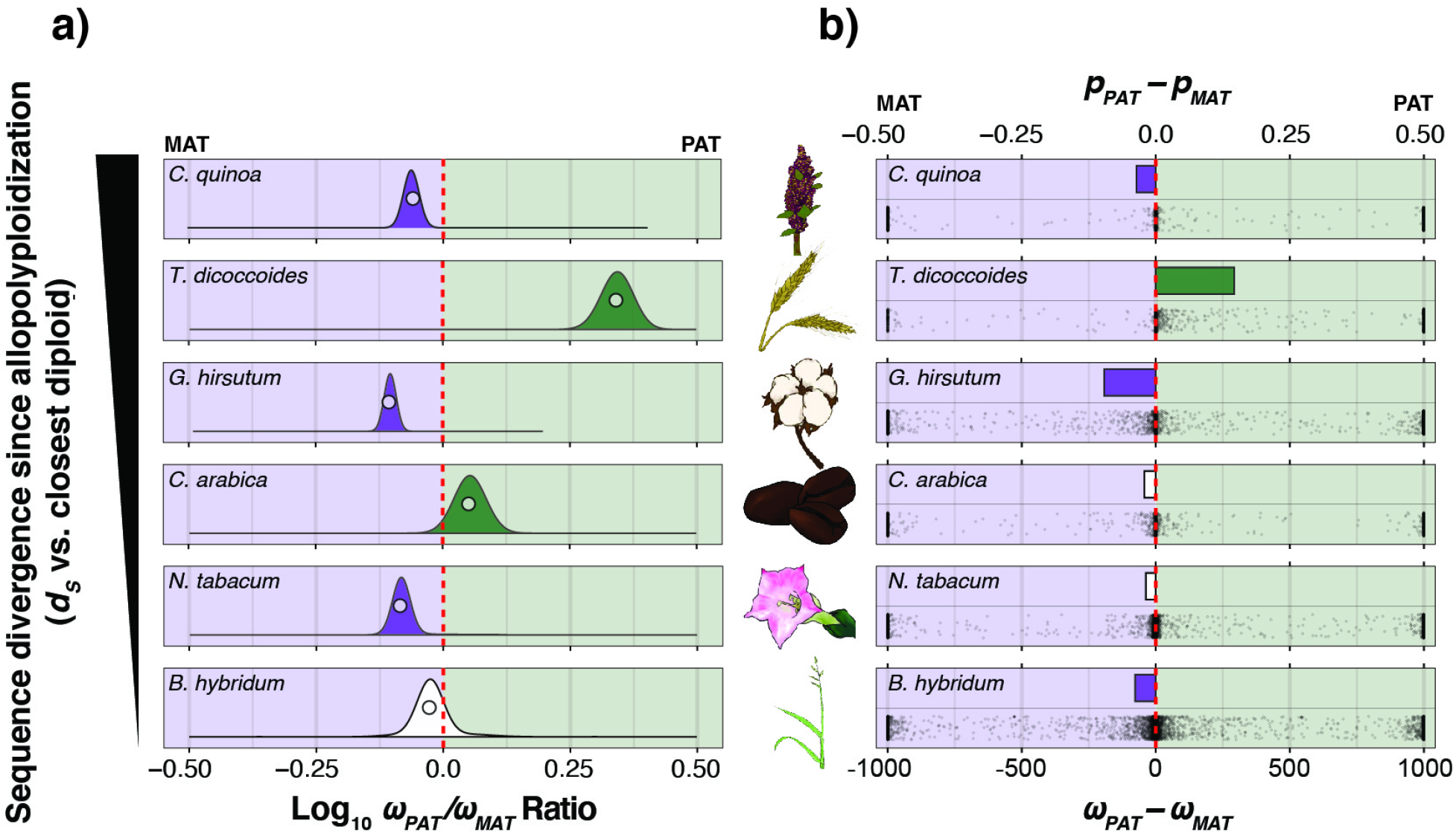
Genome-wide bias in *ω* (*d_N_/d_S_*) across maternal and paternal subgenomes. a) Log-transformed ratios of *ω* values in paternal (*ω_PAT_*) vs. maternal (*ω_MAT_*) subgenomes from concatenations (circles), and underlying bootstrap distributions (density curves) of genes encoding proteins that are not targeted to either the plastids or mitochondria. Species panels are arranged vertically from oldest (top) to youngest (bottom). The red-dashed line indicates equal *ω* values across subgenomes, left of the red line indicates higher *ω* values in the maternal subgenomes, and right of the red line indicates higher *ω* values in the paternal subgenome. Bootstrap distributions of *ω* ratios that depart significantly (*p* < 0.05) from the red line are filled in solid according to the direction of subgenomic bias (i.e., green: *ω_PAT_*/*ω_MAT_* > 1.0; purple: *ω_PAT_*/*ω_MAT_* < 1.0; no fill: *ω_PAT_*/*ω_MAT_* ≈ 1.0). b) Estimates of *ω_PAT_* – *ω_MAT_* for each individual gene is depicted on the bottom half of each species’ panel and the proportion of genes with higher *ω* values in the paternal subgenome (*p_PAT_*) minus the proportion of genes with higher *ω* values in the maternal subgenome (*p_MAT_*) is depicted on the top half of each species’ panel for all genes not-targeted to either the mitochondria or plastids. The red-dashed line represents equal proportions of genes with higher *ω* values across subgenomes, and bars are filled in when proportion deltas are significantly different from zero (i.e., green: *p_PAT_* > *p_MAT_*; purple: *p_PAT_* < *p_MAT_*; no fill: *r_PAT_* ≈ *r_MAT_*).

We next performed concatenated and gene-level analyses of *ω_PAT_* and *ω_MAT_* in organelle-targeted genes (normalized by NOT genes) to test whether paternal homoeologs exhibited faster rates of protein-sequence evolution than maternal homoeologs, as predicted if paternal subgenomes harbor incompatibilities with the cytoplasmic genomes. We found evidence that concatenations of wheat genes involved in mitochondrial enzyme complexes exhibited significantly higher *ω_PAT_* values (median = 0.661, 95% CI = 0.268 – 0.807) compared to *ω_MAT_* values (median = 0.0771, 95% CI = 0.0460 – 0.125), relative to NOT genes (*ω_PAT_* = 0.444, 95% CI = 0.414 – 0.476; *ω_MAT_* = 0.201, 95% CI = 0.189 – 0.215); however, no other species or functional classes exhibited the predicted pattern (Figure 5). To further investigate the patterns of molecular evolution the wheat mitochondrial enzyme complex genes, we manually inspected and trimmed concatenated alignments from NOT genes, mitochondrial enzyme complex genes, and plastid enzyme complex genes and re-inferred *ω_PAT_* and *ω_MAT_* in all three gene categories. Importantly, we found two small regions from two genes in the mitochondrial enzyme complexes that were poorly aligned only in the paternal subgenome, contributing to elevated *ω_PAT_* but not *ω_MAT_*. The poorly aligned regions appeared to be caused by a combination of an apparent frameshift in the paternal homoeologs of one gene encoding a protein involved in the NADH Dehydrogenase (OXPHOS Complex I – TRIDC1AG048530) and another gene encoding a protein that functions in the large subunit of the mitoribosome (TRIDC4AG029590) had an exon on the 3’ end of the gene with no apparent homology to the other sequences in the quintet (likely due to misannotation or misassembly, as the new *T. turgidum* assembly, GCA_900231445.1, does not have this same issue). Both genes exhibited substantially different *d_S_* and *d_N_* values compared to other genes in the same functional gene category (Table S4). Trimming the poorly aligned regions resulted in a substantially lower *d_N_* value for concatenated alignments of mitochondrial enzyme complex genes, which in turn caused a lower *ω_PAT_* value that was not significantly different from the *ω_MAT_* value (Figure S6). All trimmed alignments and analyses are available at https://github.com/jsharbrough/allopolyploidCytonuclearEvolutionaryRate. For gene-level analyses, we did not find any functional categories in any species that exhibited significantly different normalized proportions of genes with higher *ω_PAT_* or *ω_MAT_* (Figure S7), a pattern which did not change when *d_N_* was used in place of *ω*. Thus, there do not appear to be global accelerations in protein-sequence evolutionary rate of paternal homoeologs of organelle-targeted genes in the wake of allopolyploidization.

**Figure 5.**
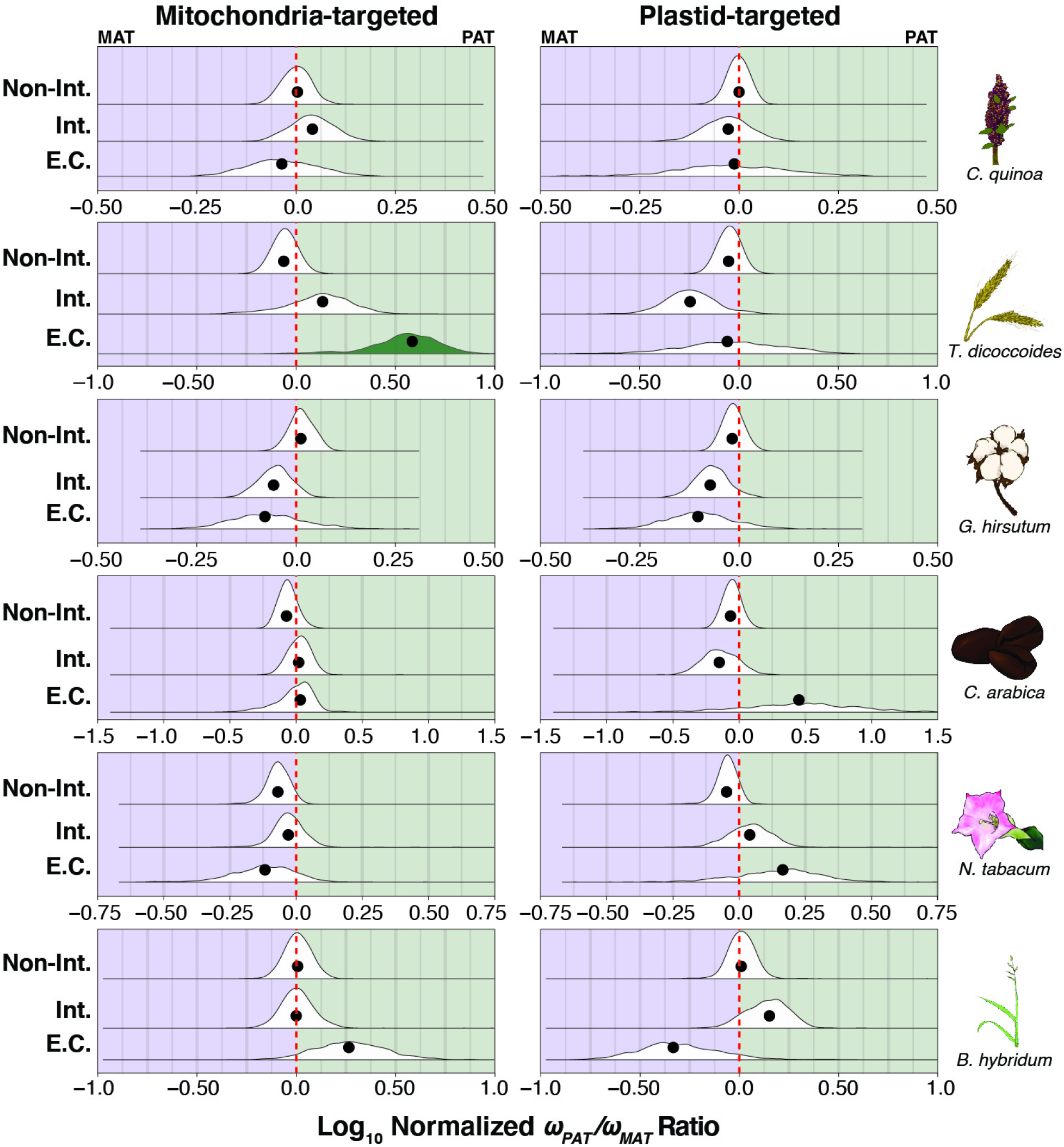
Ratios of maternal vs. paternal *ω* values in organelle-targeted genes. Log-transformed ratios of maternal vs. paternal *ω* values for concatenations (black circles) and underlying bootstrap distributions (density curves) of mitochondria- (left) and plastid-targeted (right) genes in six focal allotetraploid species. Species panels are arranged vertically from oldest (top) to youngest (bottom). The red-dashed line indicates the *ω_PAT_*/*ω_MAT_* ratio for a concatenation of genes not-targeted to the organelles (Figure 4a). Ratios left of the red line indicate higher *ω* values in the maternal subgenome, and ratios right of the red line indicate higher *ω* values in the paternal subgenome, after accounting for genome-wide patterns. Bootstrap distributions of *ω* ratios that depart significantly (*p* < 0.05) from the red line are filled in solid according to the direction of subgenomic bias (i.e., green: normalized *ω_PAT_*/*ω_MAT_* > 1.0; purple: normalized *ω_PAT_*/*ω_MAT_* < 1.0; no fill: normalized *ω_PAT_*/*ω_MAT_* ≈ 1.0). The intimacy of interactions are indicated on the y-axis from low or no interaction with organelle gene products (top), to interacting genes (middle), to genes involved in mitochondrial or plastid enzyme complexes (bottom).

We next evaluated *ω* values at the level of specific cytonuclear interactions (Table S5) and found scattered patterns of both paternal and maternal bias across various cytonuclear interactions in the three older polyploids (i.e., quinoa, wheat, and cotton). In particular, paternal homoeologs of quinoa exhibited significantly higher *ω* values (i.e., *ω* values from concatenated alignments +/- 1 SE were outside bootstrap-constructed 95% confidence intervals of NOT genes) than maternal homoeologs in mitochondrial tRNA base modification, plastid NDH, and plastid tRNA base modification, and maternal homoeologs exhibited significantly higher *ω* values than paternal homoeologs in both subunits of the chlororibosome and Photosystem I (PSI). As seen at higher levels of organization, wheat mitochondrial enzyme complexes generally exhibited higher *ω* values in paternal vs. maternal homoeologs (see below for detailed discussion) compared to NOT genes. However, the reverse was true in plastid enzyme complexes, with plastid PSII exhibiting significantly higher *ω* values in maternal vs. paternal homoeologs, relative to NOT genes. Wheat organellar tRNA aminoacyl synthetases, which are largely dual-targeted [125], also exhibited significant maternal bias compared to NOT genes. Cotton had fewer CyMIRA categories that showed evidence of bias over-and-above genome-wide levels, with just the mitochondria- and plastid-targeted recombination, replication, and repair (RRR) genes (also commonly dual-targeted [67]) exhibiting elevated *ω* values in paternal vs. maternal homoeologs and the large subunit of the mitoribosome and mitochondria-targeted PPR genes exhibiting higher *ω* values in maternal vs. paternal homoeologs compared to NOT genes. Coffee, tobacco, and *Brachypodium* all appear to be too young for this analysis, as only a single functional category (plastid transcription and transcript maturation) in coffee showed significant (maternal) bias compared to NOT genes, despite genome-wide bias in *ω* values of coffee and tobacco. There were no CyMIRA categories that exhibited consistent patterns across even the older three allopolyploids, highlighting the highly context-specific nature of evolutionary dynamics of cytonuclear interactions in allopolyploids.

Because incompatibilities are only likely to arise in genes that are divergent at the time of allopolyploidization, we also performed the analyses described above on high and low divergence gene bins. To do so we split single-copy orthologous quintets into two groups: those with high amino acid sequence divergence between diploid models (measured by *d_N_*) and those with low amino acid sequence divergence. We used a similar approach as before to normalize *ω_PAT_* and *ω_MAT_* using the NOT genes. There were only two cases in which high and low divergence classes differed by more than one standard error: mitochondrial and plastid enzyme complexes of wheat (Figure S8). In particular, the low divergence class of wheat mitochondrial enzyme complexes (MTEC) exhibited more extreme paternal bias than the high divergence class, while the low-divergence class of wheat plastid enzyme complexes (PTEC) exhibited a more extreme maternal bias compared to the high-divergence class. This somewhat surprising result notwithstanding, the lack of signal in the high divergence classes across the other functional categories and species indicates that cytonuclear incompatibilities of allopolyploids are not resolved by faster rates of protein-sequence evolution in paternal homoeologs.

We compared patterns of autapomorphic amino acid changing mutations at sites that were conserved throughout the rest of the quintet in genes encoding subunits of mitochondrial enzyme complexes. For each species, we observed several gene functional categories that exhibited an excess number of autapomorphic amino acid changes compared to genes not targeted to the mitochondria or plastids in one subgenome compared to the other. However, the direction of excess was not consistent across species or even across functional gene categories (Table S6).

Because derived amino acids with substantially different biochemical properties compared to ancestral residues (i.e., radical amino acid changes) are especially likely to alter protein structure and function [126–131], we next restricted these analyses of derived amino acid changes in the tetraploids to only include radical amino acid changes (as defined by the Conservative/Radical Index CRI [132]). As was the case with total derived amino acid changes, there existed several functional categories in each species that exhibited significant biases in the accumulation of radical autapomorphies across subgenomes, but the direction of bias and the functional categories identified were not consistent across species. Several notable functional categories did exhibit bias across multiple species though (e.g., DNA replication, recombination, and repair genes [quinoa, cotton, *Brachypodium*], tRNA base modification genes [quinoa, cotton, coffee, *Brachypodium*], and tRNA aminoacyl synthetases [wheat, tobacco]), potentially indicating they are hotbeds for cytonuclear incompatibilities and/or diploidization. Together these results indicate that cytonuclear enzymes exhibit complex- and species-specific patterns of accumulation of derived amino acids at conserved sites.

In sum, our concatenated, gene-level, and site level analyses provide evidence that protein sequences of different allopolyploid subgenomes exhibit different *ω* values, potentially as a result of different rates of protein-sequence evolution, but cytonuclear incompatibilities resulting from the allopolyploidization event do not leave global signatures of accelerated protein sequence evolution in paternal homoeologs of organelle-targeted genes. Moreover, while organelle-targeted genes are often lost at higher rates than genome-wide rates of diploidization, this is not always the case, especially in cotton, and biased gene content of allopolyploid subgenomes does not appear to be related to cytonuclear incompatibilities. Rather, only species- and complex-specific cytonuclear dynamics appear to alter rates of evolution in organelle-targeted genes, and in directions not uniformly consistent with allopolyploidy induced cytonuclear incompatibilities.

## DISCUSSION

We inferred orthologous gene sets, partitioned genes by subcellular targeting localization and cytonuclear interaction, and evaluated genome-wide patterns of gene content and natural selection across subgenomes of six allotetraploid angiosperms. We report significant genome-wide biases across maternal vs. paternal subgenomes in overall gene content in all five allopolyploids tested and in mutation-rate-corrected rates of protein-sequence evolution (i.e., *ω*) in all six allopolyploid genomes tested. The directions of bias in both gene content and higher *ω* were not consistent across independent allopolyploidization events, and the patterns observed in gene content did not appear to be similar in direction as bias in *ω*.

The analyses reported here support three primary conclusions: (1) allopolyploid subgenomes exhibit substantially different rates of protein-sequence evolution from one another despite existing inside the same cell for thousands to millions of years; (2) cytonuclear incompatibilities between the cytoplasmic genomes and the paternal subgenome are complex and taxon-specific and do not result in global increases in rates of protein-sequence evolution in paternal homoeologs of organelle-targeted genes; and (3) gene content is not equally distributed across subgenomes, with both species and cytonuclear functional classes contributing to variation in the rate at which genomes fractionate following WGDs. The foregoing conclusions suggest a number of questions that have implications for our understanding of polyploid biology.

### Differential rates of protein-sequence evolution across allopolyploid subgenomes

Most prominent among our data are the remarkable differences in evolutionary patterns across subgenomes, raising the question of what evolutionary forces underlie these subgenomic biases? That is, allopolyploid subgenomes that have been (co-)evolving inside the same nucleus for thousands to millions of years [133], remain on separate evolutionary trajectories with respect to evolutionary rates in protein-coding genes. Here, we consider several phenomena that could play a role in establishing and maintaining subgenomic biases.

If *ω* is adequately inferring patterns of natural selection across subgenomes (but see below for alternative explanations), then the patterns of subgenomic biases in rates of protein-sequence evolution reported here could arise from differences in the efficacy of selection or effective population size (*N_e_*) across subgenomes. In particular, genes that are more highly expressed [134, 135], have higher local recombination rates [136–139], or lower local TE densities [140–142] (but see [143]), are expected to experience increased efficacies of natural selection and thus exhibit reduced rates of protein-sequence evolution [144]. That is, genome-wide differences between the progenitors at the time of allopolyploid formation (e.g., transcriptome size, recombination rate, TE load) would not only be expected to give rise to subgenomic differences in the immediate aftermath of polyploidization [101,145–148], but also contribute to evolved differences across subgenomes [10,11,149–153].

Mutation rate varies tremendously across species, populations, individuals, and even within genomes [154–157], making it a potential candidate for generating subgenome biases in *ω*, if elevated mutation rate results in increased rates of background selection, thereby reducing *N_e_* [144]. Such mutational biases could reflect ancestral differences in parental species (e.g., differences in DNA methylation [157]), or could potentially arise after polyploidization in association with other biased phenomena such as recombination [158], gene expression [8,36,38,159–166], epigenetic marks [26–33], or transposable element activity [14–17], which are all thought to themselves be mutagenic [167–170].

Subgenomes might also differ in *N_e_* as a result of backcrossing, in which one polyploid subgenome experiences higher rates of introgression than the other [171–173]. Repeated allopolyploid formation or gene flow from diploids (e.g., *Brachypodium hybridum* – [122], *Arabidopsis suecica* – [174]) can cause *N_e_* to differ across subgenomes. Finally, recombination could also act to bias inferences of *ω* artifactually because genetic material exchanged across subgenomes via homoeologous exchange [18–25,175–183], gene conversion [19,20,57,62,133,153,184–189], and other recombinational mechanisms (e.g., [190]) would be expected to bias *ω* inferred across a topologically constrained tree. However, we took steps to prevent this type of artifact from influencing our data by only including genes that exhibited gene-tree topologies that were consistent with the species tree topology.

The relative contributions of these various evolutionary dynamics are of central importance to the understanding of polyploid genomes, but testing each hypothesis in turn is made difficult by the fact that the sampled diploids are, to varying degrees, imprecise models of the ancestral progenitors. Therefore, an unknown fraction of each terminal “polyploid” branch in our quintet trees actually represents evolution as a diploid prior to hybridization. Wheat in particular is susceptible to artifactual inflation of *ω* because *Aegilops speltoides* is so much more distantly related to the B subgenome of the polyploid than *Triticum urartu* is to the A subgenome (Figure 2). The persistence of slightly deleterious changes since the divergence of the A subgenome and the diploid A genome may result in overestimates of *ω* in the A subgenome compared to the B subgenome. The same logic applies to all of our polyploid taxa to varying extents; however, it is worth noting that while differences in *d_S_* across subgenomes were the primary drivers of differences in *ω* in wheat and tobacco, *d_N_* had a proportionally larger effect than *d_S_* on differences in *ω* in quinoa, cotton, coffee, and *Brachypodium*. This latter finding is consistent with selection being the driving factor in variation in evolutionary rates across subgenomes (but see prior caveat regarding quality of diploid models and evolution prior to polyploidization), rather than mutation rate variation or artifactual inflation of *ω* in the more closely related diploid-subgenome pair. In the same vein, coffee and cotton, which are both thought to have extremely small effective population sizes [191], exhibited the highest overall *ω* values (Figure S5b). All told, investigating site frequency spectra, gene expression profiling, and recombination rates within populations and their relationships to the biased *ω* values reported here will help resolve these outstanding questions.

### No global signature of mitonuclear incompatibilities in paternal homoeologs of allopolyploid genomes

To test the hypothesis that incompatibilities stemming from evolutionary mismatches between the maternally derived cytoplasmic genomes and the paternally derived nuclear subgenome result in preferential loss and accelerated rates of protein-sequence evolution in paternal homoeologs of organelle-targeted genes, we applied the same analyses described above to sets of CyMIRA-partitioned genes, after accounting for genome-wide effects. We did not discover evidence that cytonuclear incompatibilities shape either gene content or protein-sequence evolution in paternal homoeologs of organelle-targeted genes, despite multiple distinct tests of this hypothesis. In particular, patterns of gene content on organelle-targeted genes exhibited the opposite pattern as that observed in NOT genes in three of five allopolyploid taxa (the remaining two were not significantly different from genome-wide patterns), indicating that organelle-targeted genes tend to exhibit greater balance across subgenomes than the rest of the genome. While the proportion of organelle-targeted genes per subgenome did not appear to be especially maternally biased, four of six allotetraploids had reduced overall proportions of organelle-targeted genes compared to NOT genes. Overall, rates of protein-sequence evolution in organelle-targeted and interacting genes generally reflected the genome-wide patterns of bias observed in NOT genes, rather than rate accelerations peculiar to paternal but not maternal homoeologs.

One outstanding question stemming from our analyses of protein-sequence evolution in paternal vs. maternal homoeologs of organelle-targeted genes is why hybrid polyploid genomes appear to generally lack genome-wide signatures of cytonuclear incompatibilities, despite their apparent importance in homoploid hybridization [95] and introgression events [92]? It is possible that cytonuclear incompatibilities do leave signatures on genomes, but not in terms of accelerated rates of protein-sequence evolution in paternal homoeologs. For example, pseudogenization may be a rapid and common mechanism for adaptation in plant genomes (e.g., [192]), which would be missed by our quintet analyses. While we did not observe maternally biased gene content in CyMIRA datasets, direct analysis of bias in homoeologous pairs in which one copy is pseudogenized is necessary to rule out gene loss as a mechanism by which cytonuclear incompatibilities are resolved. The seemingly stochastic patterns of homoeolog bias in accumulation of autapomorphic amino acid changes indicates that there are often cases in which homoeologs of cytonuclear interacting genes evolve very differently, perhaps reflecting cytonuclear incompatibilities or the precursor to gene loss and diploidization, but these biases do not appear to coincide with the allopolyploidization events in any systematic way. The presence of biased accumulation of autapomorphies in *Brachypodium* may indicate that cytonuclear incompatibilities are resolved rapidly. Cytonuclear incompatibilities may also be resolved via biased homoeolog expression [162], gene conversion [57, 62] or homoeologous exchange [25], subfunctionalization of subcellular localization by differential isoform usage across homoeologs [193], or other potential mechanisms that would not generate global signatures of paternal acceleration in coding sequences of organelle-targeted quintets.

Biased homoeolog expression represents a potential mechanism by which allopolyploids could resolve cytonuclear incompatibilities, but has found mixed support in the studies that have so far attempted it. In particular, cotton, tobacco, *Arabidopsis*, peanut, and the extremely young allotetraploid *Tragopogon miscellus* exhibit biased maternal expression of the nuclear-encoded subunit of Rubisco [56–58], but others have not found similar patterns in rice [59] or *Brassica napus* [60]. Moving forward, large-scale genome-wide homoeolog expression bias could be evaluated across all the CyMIRA gene sets (not just Rubisco) to test this hypothesis. Additionally, the topological and alignment filtering steps we imposed on quintets here had the intended side effect of filtering out genes exhibiting gene conversion or homoeologous exchange. Notable among them was *rbcS*, which encodes the small subunit of Rubisco and was missing from filtered, single-copy quintets in five of six species complexes (present only in *Brachypodium*, the youngest allopolyploid). It is likely that because of *rbcS*’ propensity for gene conversion [57], this apparent “hotbed” for cytonuclear incompatibilities might provide additional evidence that was missed here. Certainly, a careful analysis of maternal vs. paternal bias in gene conversion tracts and homoeologous exchanges among organelle-targeted genes may be a fruitful future approach.

An additional and perhaps likely possibility is that cytoplasmic genomes of these allopolyploids may evolve too slowly in protein-coding sequence to generate widespread incompatibilities in hybrid polyploids [194]. The relatively young allopolyploid *Brassica napus* may be a relevant example. The plastid genomes of *B. oleracea* and *B. rapa* have very few differences, and a recent analysis did not detect extensive incompatibilities with nuclear subgenomes [60]. By contrast, elevated rates of protein-sequence evolution and *ω* values in organelle-interacting genes have been detected repeatedly in lineages with rapidly evolving cytoplasmic genomes [86,87,89,195–202]. Therefore, genome-wide analyses of evolutionary rates appear to be sensitive enough to detect cytonuclear incompatibilities when their effects are strong.

Because cytonuclear interactions are critical for hybrid lineage success in many cases [203–205], allopolyploids with cytonuclear incompatibilities may also be evolutionarily short-lived, such that the relatively successful allopolyploids assayed here may be unlikely to exhibit cytonuclear incompatibilities. Along these lines, allopolyploid unisexual salamanders do not appear to exhibit maternally biased expression of nuclear-encoded OXPHOS genes [206], despite high rates of mitochondrial DNA sequence evolution and ancient divergence of the mitochondrial lineage from the paternal lineages [173]. The high incidence of asexuality and selfing species among polyploid lineages may speak to this possibility [44]. Overall, the data presented here and elsewhere appear most consistent with a scenario in which cytonuclear incompatibilities have minimal effects on rates of protein sequence evolution in allopolyploid plants.

### Cytonuclear gene content evolution in allopolyploids

Polyploids often have both larger cells [52,207–210] and more chloroplasts per cell in leaf tissue ([211–216][53]). Together, these phenomena suggest that stoichiometry between nuclear and cytoplasmic genomes is important for cellular and organismal function [55]. Previous work investigating single-copy genes in plants indicated that organelle-targeted genes are among the first to return to diploidy following whole genome duplication events [123, 124]. The gene content analyses presented here generally agree with those analyses, although cotton and coffee offer important exceptions that muddy the waters. By contrast, Ferreira de Carvalho and colleagues [60] reported higher levels of maintained duplicates in organelle-targeted genes in the allopolyploid *Brassica napus,* compared to genome-wide levels. The discrepancies between the former two studies (performed in diploids) and the latter two (performed in polyploids) indicates that cytonuclear stoichiometry may be highly responsive to nuclear gene content. In support of that hypothesis, diverse polyploids appear to compensate for elevated nuclear ploidy with increased organelle genome copy number [40,214,217–220]. Additional work investigating the immediate and evolved consequences of cytonuclear stoichiometry at the genomic, transcriptomic, proteomic, and organellar levels, especially by homoeologous pair analysis, will provide valuable insights into the unresolved question of how genome doubling can affect cellular energy production and homeostasis.

### Summary

The genome-wide analyses of maternal vs. paternal evolutionary rates presented here represent the most extensive investigation of cytonuclear incompatibilities in allopolyploids performed to date, representing six distinct allopolyploidization events of varying ages and divergences. We find clear evidence of differential evolution across subgenomes, but little evidence of paternal-homoeolog-specific accelerations of evolutionary rates in organelle-targeted genes. Additionally, we found that organelle-targeted gene content tends to be less biased than the rest of the genome, with mixed evidence of whether organelle-targeted genes tend to be lost more often than the rest of the genome. Further work investigating the forces underlying these observations and the consequences for organismal energy metabolism and homeostasis will be critical for understanding the cytonuclear dimension of allopolyploidy.

## MATERIALS AND METHODS

### Genomic datasets

The proliferation of genome assemblies for polyploid plants and their diploid relatives has enabled powerful phylogenomic analyses. We identified six allotetraploids that share hybrid origins (Figure 1a), have publicly available chromosome-scale genome assemblies for both the polyploid and the diploids that are most closely related to each subgenome (with the exception of the wild emmer wheat [*Triticum dicoccoides*] B subgenome, whose diploid relative [*Aegilops speltoides*] only has a transcriptome available), and varying degrees of divergence between their diploid progenitors and the amount of time since allopolyploidization (Figure 1b). We also included the closest available chromosome-scale assembly for an outgroup species to polarize substitutions. Accession numbers and references are provided for assemblies and annotations used from each species complex in Table 5.

**Table 5.**
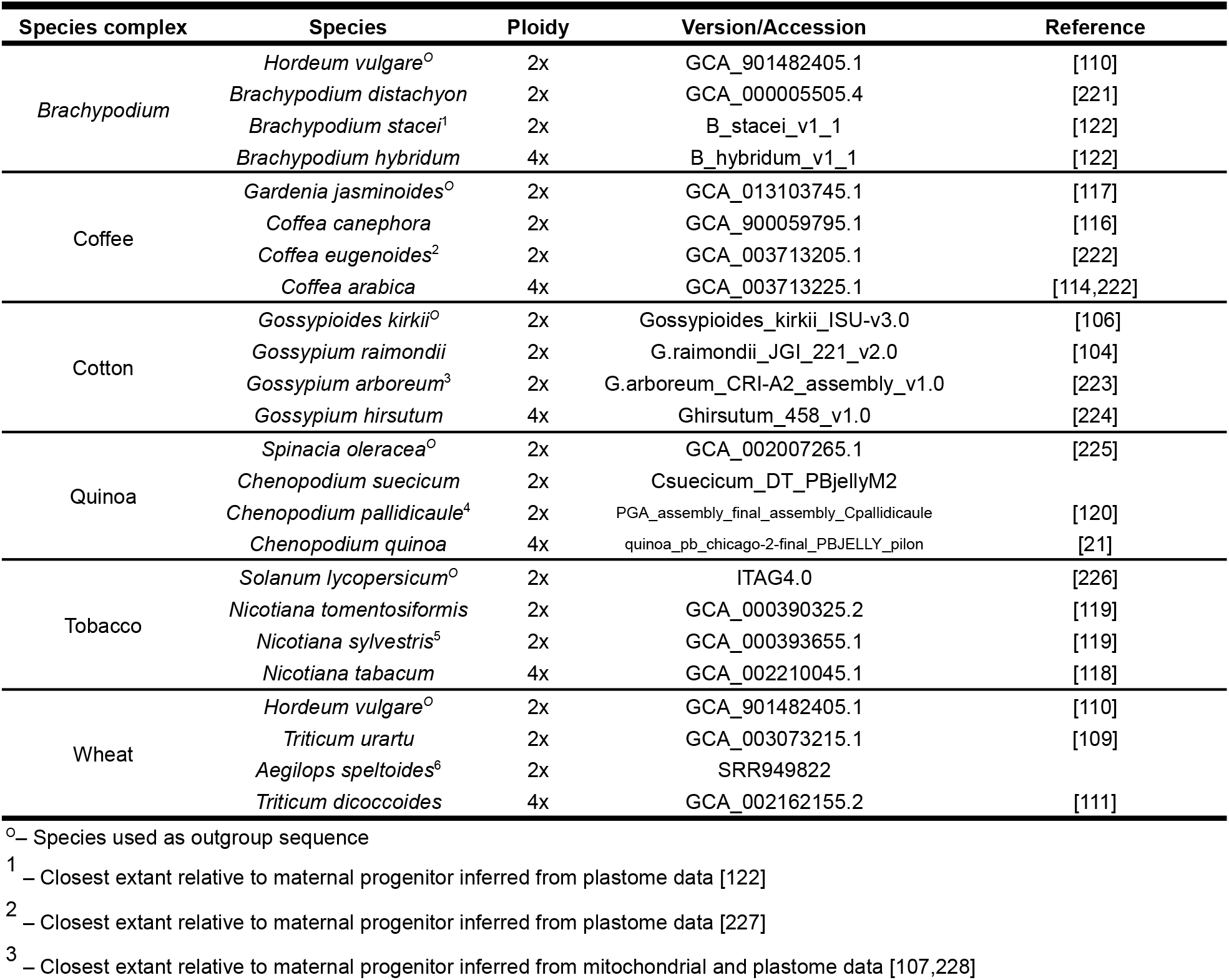
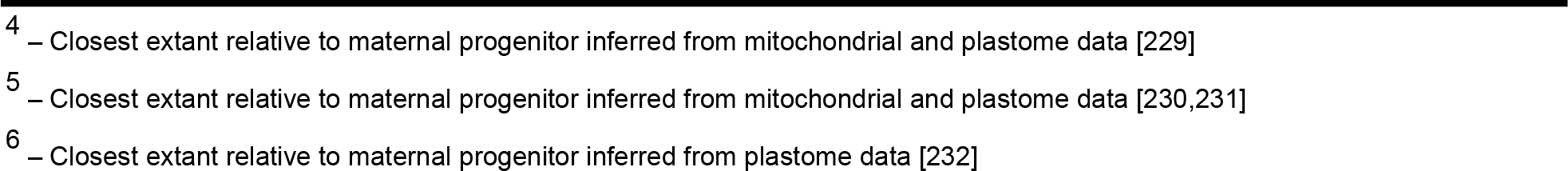
Genomic resources for six allotetraploid species complexes.

### Orthologous quintet inference

Each of the six allopolyploids have subgenomes that are more closely related to those of the sampled diploids than they are to each other. Combined with an outgroup lineage, the resulting tree topology characteristics of allopolyploids (Figure 2) allow for robust inference of lineage-specific rates of evolution in orthologous quintets. We used a combination of phylogenetic and syntenic methods to construct orthologous quintets (Figure S1).

To infer orthologous quintets using phylogenetic methods, we used Orthofinder v. 2.2.7 to infer orthologous groups of sequences, termed “orthogroups”, from the whole proteomes (primary transcripts only) of all four species [233]. For each orthogroup, we aligned CDS sequences in a codon-aware manner using the align_fasta_with_mafft_codon subroutine in the sloan.pm perl module (available at https://github.com/dbsloan/perl_modules) which translates CDS sequences into amino acid sequences, aligns those amino acid sequences with MAFFT v7.407 [234], and reverse translates the aligned amino acid positions into the CDS sequences to produce the final alignment. We selected models of molecular evolution for each alignment using jModelTest2 v2.1.10 to identify the model with the highest AICc score [235, 236], and inferred phylogenetic trees with the MPI-compatible distribution of PhyML v3.3.20180214 [235]. Five random tree starts were performed, and the treespace was further searched using a combination of nearest-neighbor-interchange subtree pruning and re-grafting. Support for trees was assessed using 100 bootstrap replicates, and splits with ≤ 50 bootstrap support were collapsed into polytomies using collapeLowSupportBranches.py (unless otherwise stated all scripts are available at https://github.com/jsharbrough/allopolyploidCytonuclearEvolutionaryRate/tree/master/scripts).

All monophyletic, minimally inclusive, species-complete subtrees were pruned out of orthogroup trees using subTreeIterator.py. We next trimmed lineage-specific gene duplicates from subtrees using trimBranches.py, which keeps only the longest sequence or a random sequence in cases where sequence length is equal across copies. The resulting trimmed subtrees that contained exactly one sequence from each diploid and two sequences from the polyploid represented our set of phylogenetic orthologous quintets. All scripts developed for reading, writing, and manipulating trees are based on the DendroPy package (https://dendropy.org/) [237].

We used the pSONIC [238]program to create a genome-wide set of syntenic orthologs. In short, pSONIC employs MCScanX [239] to create a list of pairwise syntenic blocks between all possible pairs of species in each clade, combined with orthogroups identified from OrthoFinder [233] to choose which syntenic blocks contained the highest confidence orthologs that were direct descendants of the most recent common ancestor of all species in the clade. Notably, the filtering criteria of collinear groups from our run of pSONIC differed from its formal presentation in that we did not remove collinear groups in which more genes received a “not pass” than “pass” score, and the ends of each collinear block were not trimmed as described in the manuscript describing pSONIC. These developments were made after our analyses were performed with this tool, but before the tool was submitted and reviewed for publication.

To take advantage of both inference methods, we merged phylogenetic and syntenic orthologous quintets using mergeQuintets.py to produce a high quality set of quintets that were identical across both methods (i.e. “Intersection”) and a second set of quintets that included all identical quintets plus all the phylogenetic quintets whose members were not present in the syntenic quintets and vice versa (i.e., “Union”). Results from the Intersection dataset (Supplementary File 1, Figure S10, Figure S11) did not differ in any meaningful way from the Union, so only Union results are described in the main text. Phylogenetic quintets that overlapped with but were not identical to syntenic quintets were excluded. Likewise, syntenic quintets that overlapped with but were not identical to phylogenetic quintets were also removed from our final analysis. These conflicting quintets represent a small minority of total quintets and are likely a result of the different methods by which lineage-specific duplicates are handled in the phylogenetic vs. syntenic pipelines.

For all non-conflicting orthologous quintets, we re-aligned CDS sequences as before, trimmed alignments with Gblocks v0.91b using the codon setting with the parameter -p set to n [240]. We estimated new models of molecular evolution using jModelTest2 [235, 236], and inferred phylogenetic trees as described above. We tested whether the resulting gene tree topologies agreed with the overall species tree using the quintetTopology.py script and excluded all genes with discordant tree topologies from subsequent evolutionary rate analyses.

### CyMIRA-based gene classification

To evaluate the effect of cytonuclear interactions on subgenome-specific evolutionary dynamics, we used a combination of *de novo* targeting predictions and CyMIRA [67] to partition genes into distinct functional and interaction categories. *De novo* targeting predictions were obtained from four separate targeting prediction programs: iPSORT v0.94 [241], LOCALIZER v1.0.4 [242], Predotar 1.03 [243], and TargetP v1.1b [244]. In parallel, we used Orthofinder v2.2.7 to obtain orthology information with the *Arabidopsis thaliana* Araport 11 proteome [245]. We combined the *de novo* targeting predictions with the *Arabidopsis*-inclusive orthogroups using the geneClassification.py script. Genes were classified as cytonuclear-interacting genes if they shared the same orthogroup as *Arabidopsis* genes whose products interact with mitochondrial/plastid genomes or gene products according to the CyMIRA classifications scheme [67]. Genes present in orthogroups lacking an *Arabidopsis* cytonuclear interacting gene were classified as organelle-targeted if at least one *de novo* prediction tool indicated a mitochondrial or plastid subcellular localization for the gene product and ≥50% of *Arabidopsis* genes present in the orthogroup encode products targeted to the mitochondria or plastids according to CyMIRA. Genes with evidence of dual targeting were included in both mitochondria-targeted and plastid-targeted data partitions. The resulting genome-wide targeting predictions and CyMIRA-guided classifications are available at https://github.com/jsharbrough/allopolyploidCytonuclearEvolutionaryRate/tree/master/ge neClassification and the pipeline for performing this classification is available at https://github.com/jsharbrough/CyMIRA_gene_classification. The breakdowns of gene functional categories for each genome are provided in Table 2 and Table S2.

We next evaluated whether retention of genes targeted to the organelles differs across subgenomes by comparing CyMIRA gene counts across subgenomes for five out of six polyploid genomes (*N. tabacum* was excluded from this analysis owing to the difficulty in positively identifying subgenomic ancestry for genes lacking a corresponding homoeolog). We performed binomial tests of NOT genes against expectations of equal retention, and then used *χ^2^* tests of organelle-targeted gene groups against the genome-wide patterns observed among genes not targeted to the organelles.

### Evolutionary rate comparisons

We evaluated genome-wide signatures of cytonuclear incompatibilities in organelle-targeted genes using a combination of single gene and concatenated analyses. For all single-copy quintets whose evolutionary history was consistent with the overall species tree, we removed poorly aligned quintets by estimating the total length of the tree in terms of synonymous substitutions per site (*d_S_*) using model 1 in codeml within PAML v4.9j [246]. Maximum cutoff values for *d_S_* were determined for each species complex separately and are depicted by red lines in Figure S8.

After quality filtering, we estimated *d_N_*, *d_S_*, and *ω* for individual quintets using model 1 in codeml as above, and the RateAncestor parameter set to 1. Other PAML parameters included the getSE parameter set to 1, the gamma shape parameter set to a fixed alpha of 0 (i.e., no rate variation among codons), initial omega set to 0.4, and initial kappa set to 2. For each quintet in each functional gene category, we evaluated whether the maternal vs. paternal subgenome had a higher *ω* value and a higher *d_N_*. We used *χ^2^* tests to evaluate whether individual categories differed from the pattern observed in the group of genes not targeted to the organelles. Using the inferred mutational changes from the RateAncestor output, we also evaluated whether maternal vs. paternal subgenomes had higher numbers of radical amino acid changes (i.e., substitutions between amino acids with substantially different biochemical properties) at sites that were otherwise conserved across the quintet. Substitutions were identified as radical if their score in the CRI matrix [132] was >0.5. Accumulation of derived conservative and radical amino acid changes was analyzed in a similar manner to *ω* and *d_N_* results, using a Fisher’s Exact Test to test whether there was a difference compared to genes not targeted to the organelles.

Next, we concatenated quintets according to gene functional category and estimated *ω* in maternal vs. paternal subgenomes using similar PAML parameters as before. For each PAML run, we repeated the analysis 1000 times to adequately sample the maximum likelihood plane and found median *ω* values from the replicates for each branch. We then calculated the ratio of paternal to maternal subgenome *ω* values (*ω_PAT_/ω_MAT_*), with a ratio >1.0 indicating faster rates of amino acid sequence evolution in the paternal subgenome and a ratio <1.0 indicating a faster rate of amino acid sequence evolution in the maternal subgenome. We assessed the statistical significance of the degree to which subgenomes exhibited different rates of amino acid sequence evolution by bootstrapping concatenated alignments at the gene level. For each bootstrap replicate we randomly sampled genes with replacement from the original concatenation and ran each bootstrapped alignment through five replicate runs of PAML. The median *ω* values of these five replicates were used as the bootstrap replicate values. We then found the ratio of paternal to maternal *ω* values for each bootstrap replicate and for each gene functional category to evaluate whether bootstrapped distributions departed from 1.0. To account for evolutionary forces that are not a result of cytonuclear interactions, we normalized these ratios by dividing by the paternal to maternal *ω* ratio of genes not targeted to either organelle. We inferred two-tailed *p* values directly from bootstrap distributions. For specific cytonuclear interaction categories, which are composed of only a few dozen genes or less, we manually inspected concatenated alignments, trimmed poorly aligned regions, bootstrapped alignments at the codon level using the python script bootstrapCodons.py, and performed PAML analyses with a similar approach as before.

Because cytonuclear incompatibilities are only expected when there exists divergence between the two progenitor genomes, we also binned our quintets based on high vs. low divergence between diploids for each species and repeated the gene-level bootstrap procedure described above. First, we estimated *d_N_* between diploid relatives for each quintet individually from the gene-specific PAML runs described above and placed genes according to *d_N_* into two equally sized bins. We then tested whether genes with high levels of amino acid divergence exhibit greater accelerations in *ω* in paternal copies than in genes with lower levels of amino acid sequence divergence. We evaluated statistical significance by bootstrapping alignments at the gene level and comparing paternal to maternal *ω* ratio distributions from the same gene categories to one another.

## SUPPLEMENTAL FIGURES AND FIGURE LEGENDS

**Figure S1.**
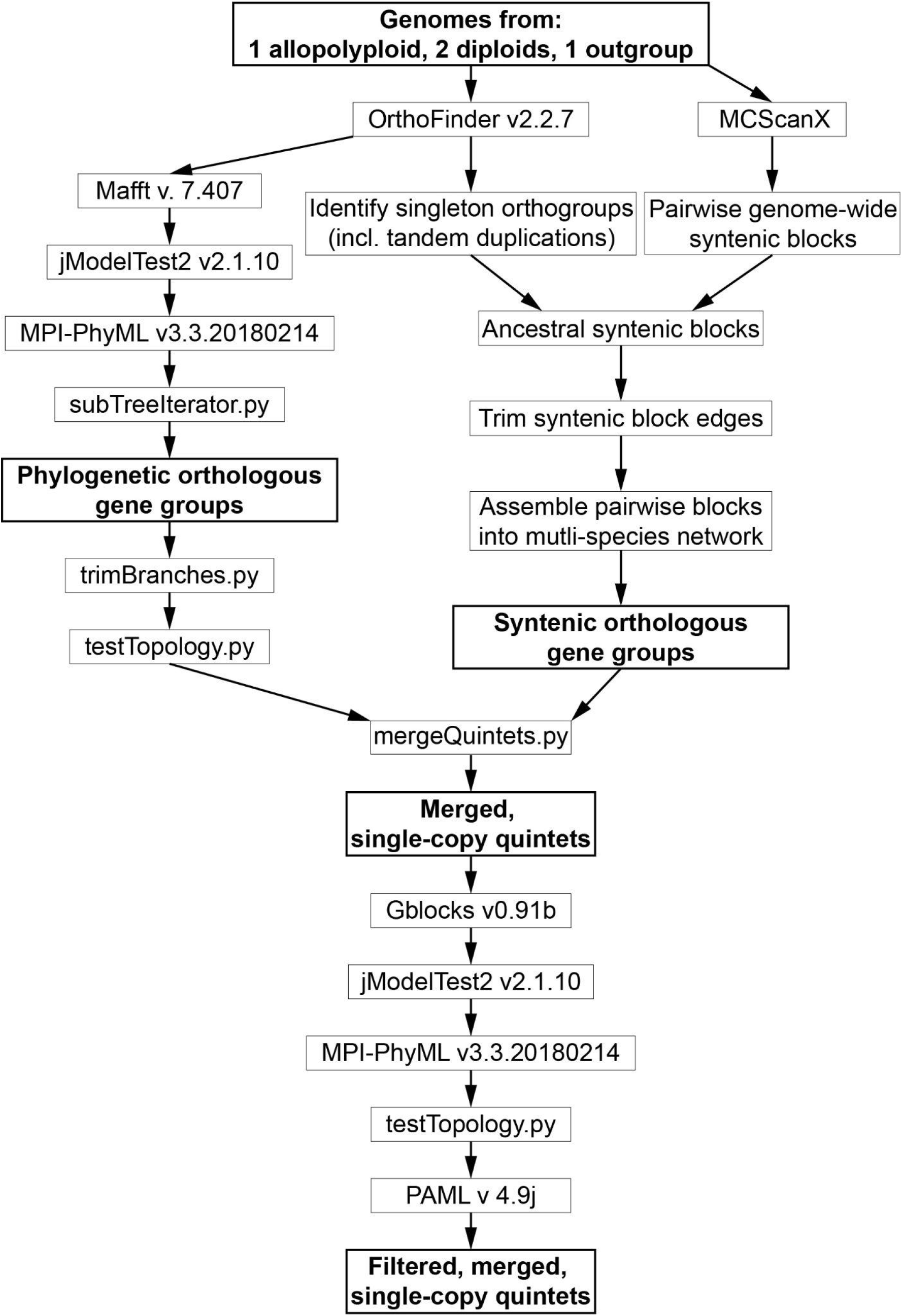
Schematic representation of phylogenetic and syntenic pipelines for inferring orthologous quintets in allopolyploid genomes. The final output of the pipeline was a set of filtered, merged, single-copy quintets, which was used in downstream analyses of rates of protein-sequence evolution.

**Figure S2.**
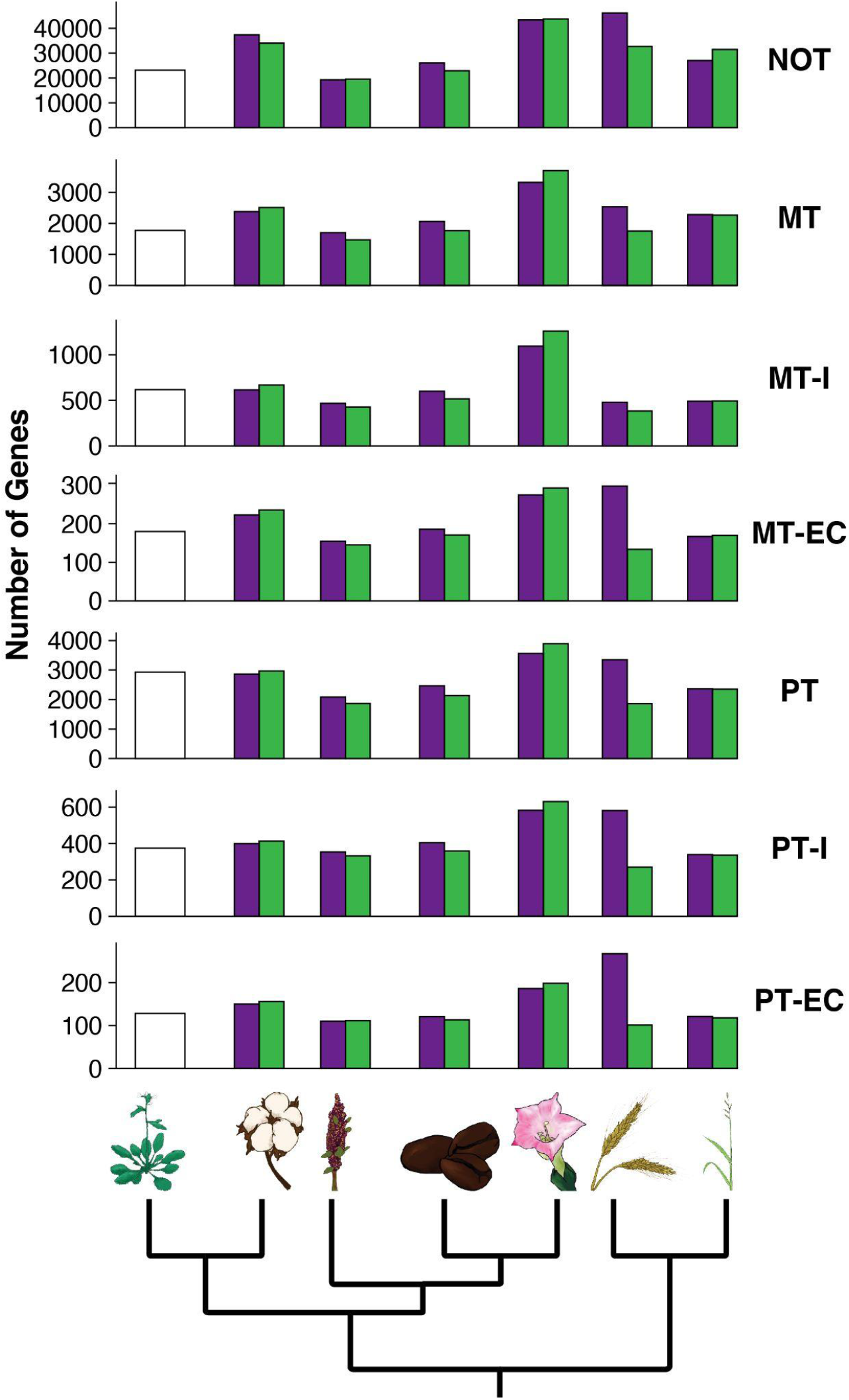
CyMIRA gene counts in diploid models compared to *Arabidopsis*. The number of genes per category is depicted on the y-axis for both maternal (purple) and paternal (green) diploid models compared to *Arabidopsis* (white). Functional gene categories are listed to the right of each plot: NOT – genes that are not-organelle-targeted, MTNI – mitochondria-targeted non-interacting genes, MTI – mitochondria-targeted interacting genes, MTEC – genes involved in mitochondrial enzyme complexes (subset of MTI), PTNI – plastid-targeted non-interacting genes, PTI – plastid-targeted interacting genes, PTEC – genes involved in plastid enzyme complexes (subset of PTI).

**Figure S3.**
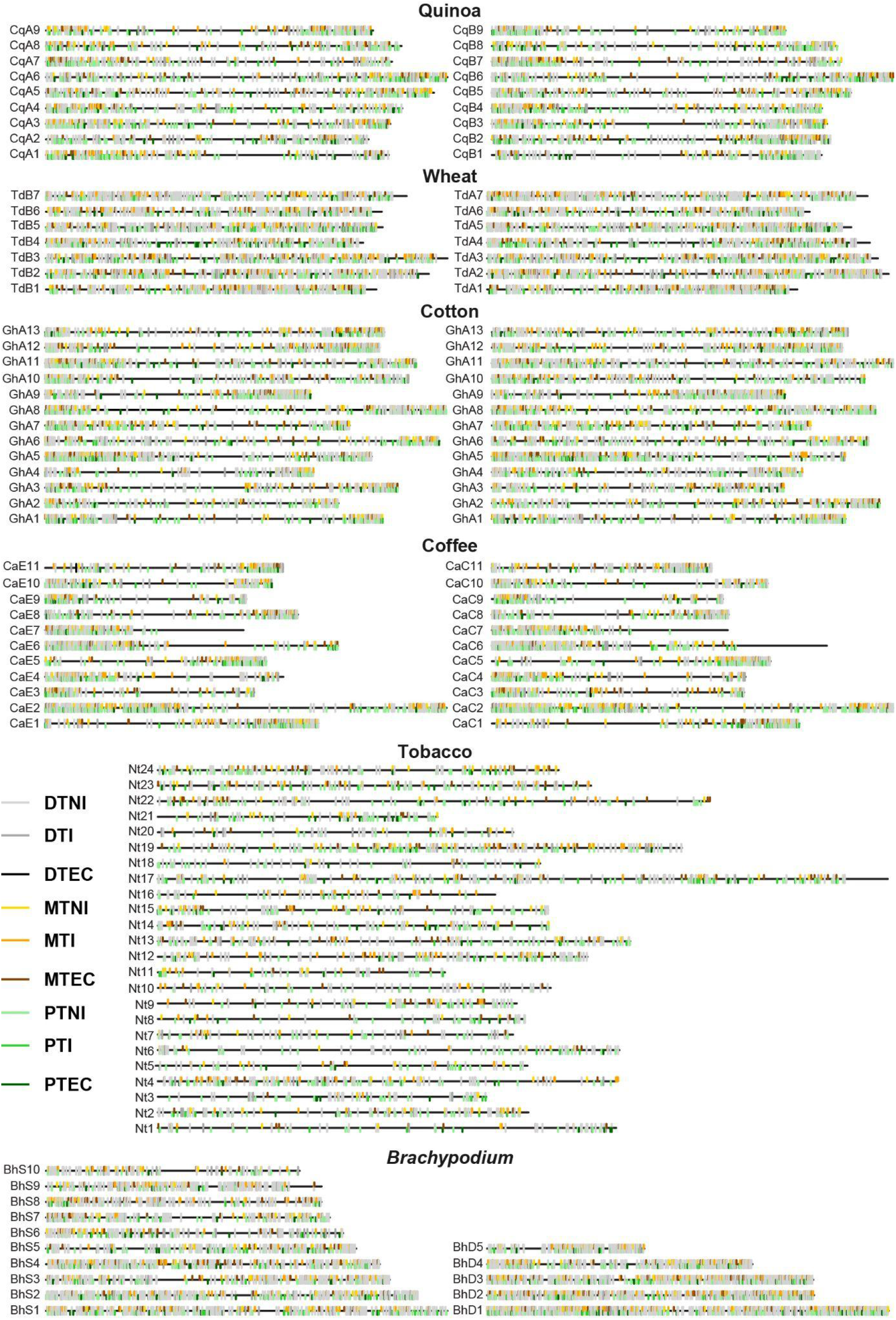
Physical distribution of organelle-targeted genes on chromosomes of focal allopolyploid genomes. Mitochondria-targeted (orange), plastid-targeted (green), and dual-targeted (grey) genes mapped onto chromosomes (black lines) of the six focal allotetraploid genomes. Taxa are arranged from oldest (top) to youngest (bottom), with maternally derived subgenomes on the left and paternally derived subgenomes on the right (excepting tobacco). Chromosome numbers are listed to the left of each chromosome (quinoa chromosomes are numbered according to similarity with *Chenopodium pallidicaule* chromosomes).

**Figure S4.**
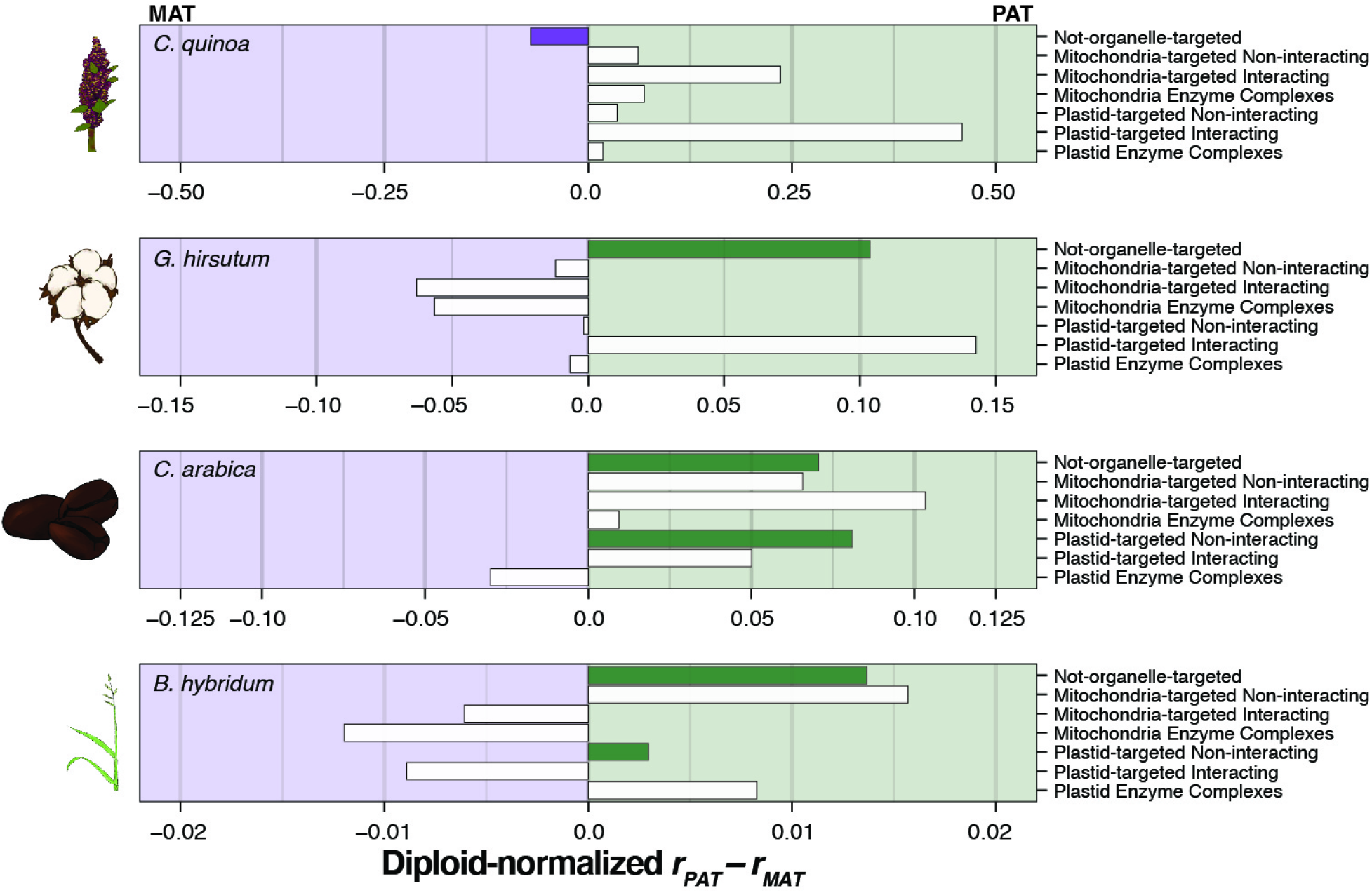
CyMIRA gene counts in maternal and paternal subgenomes of allotetraploids relative to diploid models. Bar graph depicting the number of genes present in the maternal subgenome as a proportion of the number of genes present in the maternal diploid model’s genome (*r_MAT_*) subtracted from the number of genes present in the paternal subgenome as a proportion of the number of genes present in the paternal diploid model’s genome (*r_PAT_*) for seven functional categories of genes: not-organelle-targeted, mitochondria-targeted non-interacting, mitochondria-targeted-interacting, mitochondria enzyme complexes, plastid-targeted non-interacting, plastid-targeted interacting, plasti enzyme complexes. Polyploid taxa are arranged vertically from oldest (top) to youngest (bottom) for quinoa, cotton, coffee, and *Brachypodium*. Wheat was excluded because the maternal diploid transcriptome from *Aegilops speltoides* was not a good indicator of gene counts.

**Figure S5.**
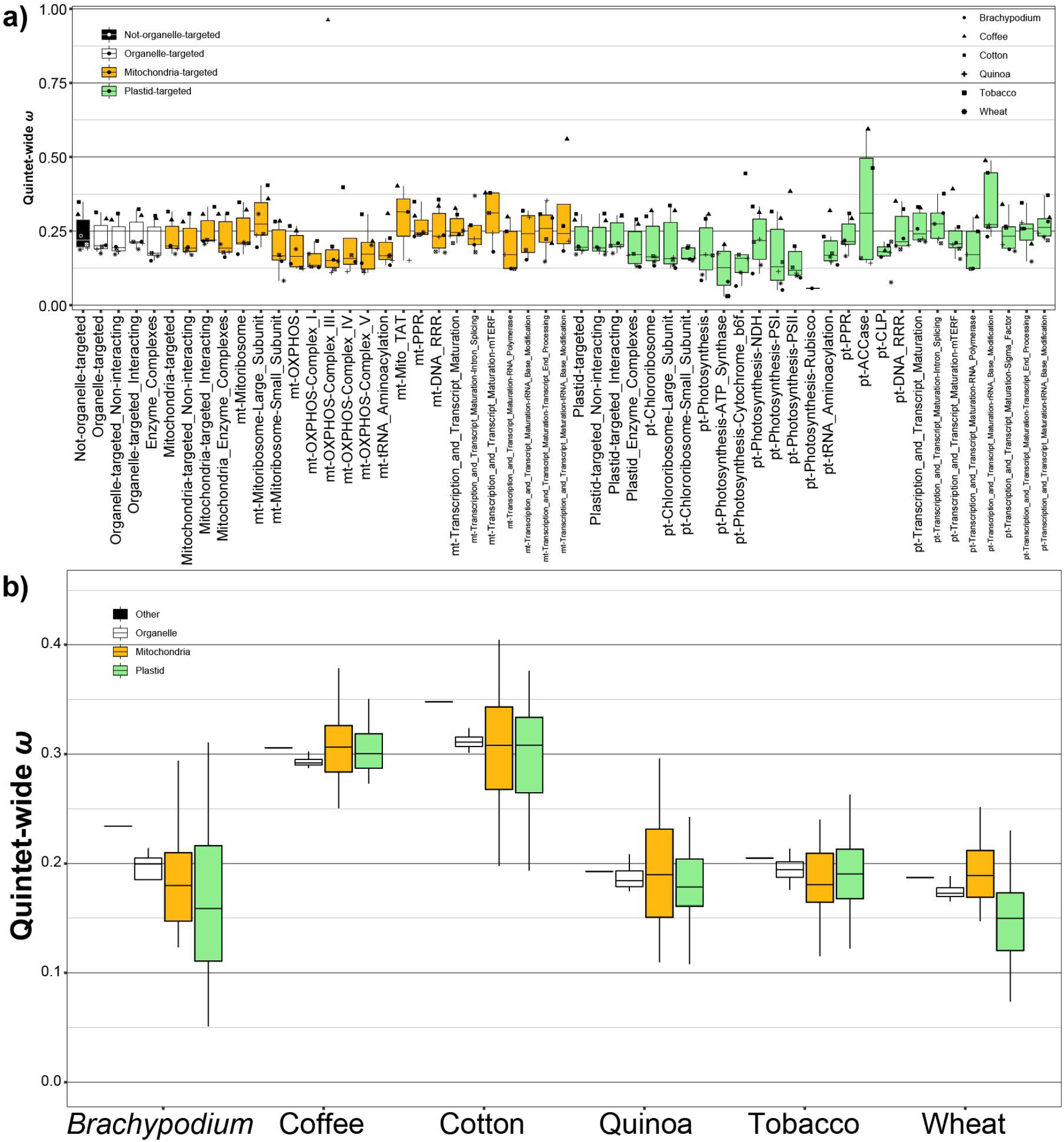
Rates of protein-sequence evolution in CyMIRA gene categories across the six focal allopolyploids. a) Quintet-wide rates of protein-sequence evolution across CyMIRA functional categories are depicted in box-and-whisker plots for the combined set of allopolyploids. Data points from each species complex are also shown, the legend for which is provided in the upper right corner of the plot. Boxes are filled according to subcellular compartment, with genes not targeted to the organelles represented by black boxes, genes targeted to either organelle by white boxes, mitochondria-targeted genes are represented by orange boxes, and plastid targeted genes are represented by green boxes. b) Boxplot depicting rates of protein-sequence evolution across allopolyploid species complexes separated by subcellular compartment of localization. Fill color is as described in panel (a).

**Figure S6.**
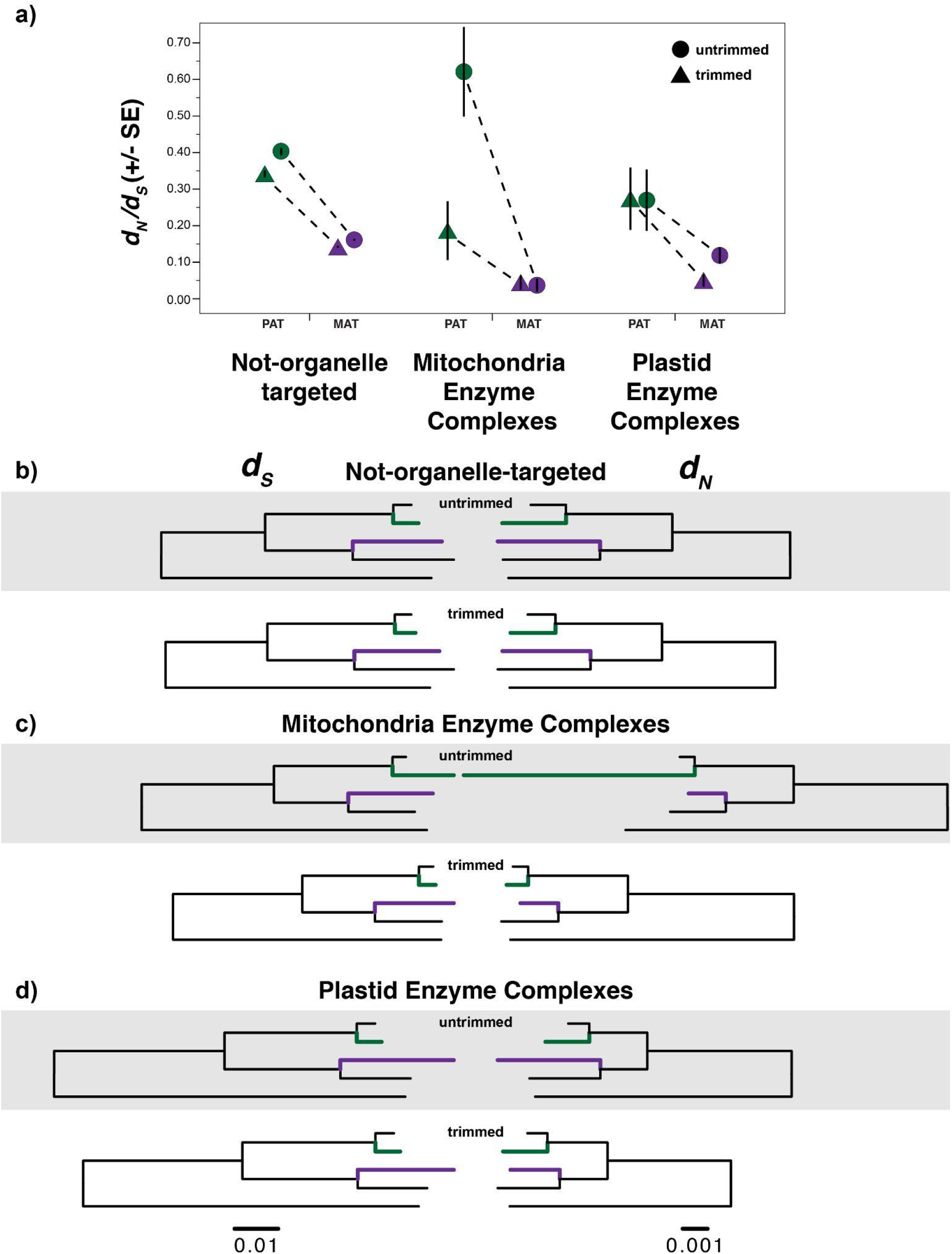
Poorly aligned regions largely explain elevated *ω* values in paternal homoeologs of wheat mitochondrial enzyme complexes. a) *ω* values from maternal (purple) vs. paternal (green) branches estimated from concatenations of genes that are not-organelle-targeted (left), involved in the mitochondrial enzyme complexes (middle), or involved in plastid enzyme complexes (right) in untrimmed (circles) vs. trimmed (triangles) alignments. The removal of two regions totalling ∼240bp accounts for the apparently elevated *ω* values in paternal homoeologs of wheat mitochondrial enzyme complex genes. b-d) Deconstructed *ω* values from concatenated PAML runs for genes (b) not-targeted to the organelles, (c) genes involved in the mitochondrial enzyme complexes, and (d) genes involved in plastid enzyme complexes in untrimmed (top) vs. trimmed (bottom) alignments. Rates of evolution for synonymous (*d_S_* - left) and nonsynonymous (*d_N_* - right) sites are represented by branch lengths, and branches are scaled similarly across functional categories. Green branches represent rates in the paternal subgenome and purple branches represent evolutionary rates in the maternal subgenome of *T. dicoccoides*.

**Figure S7.**
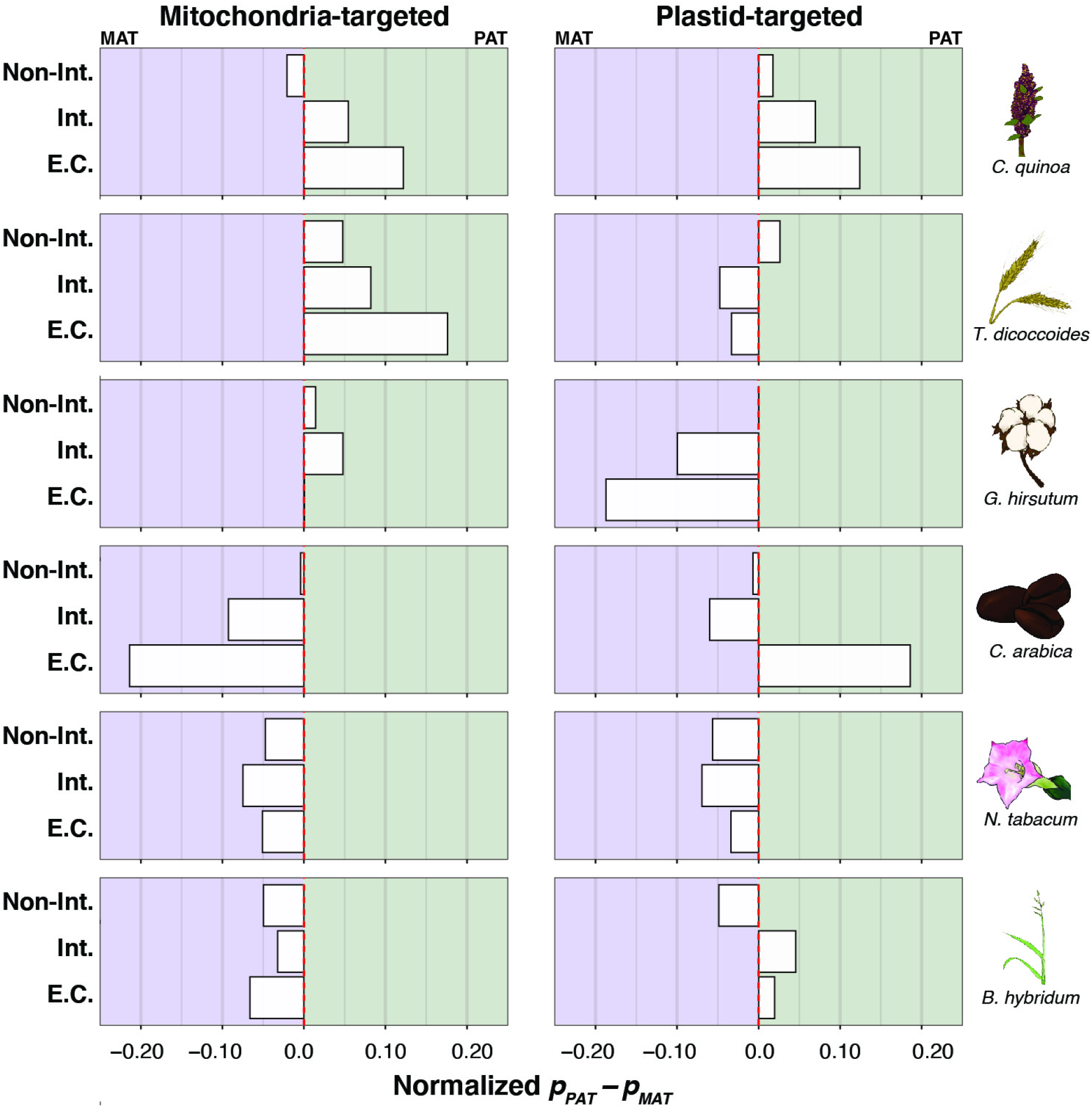
Proportions of genes that have higher *ω* values in paternal vs. maternal copies of organelle-targeted genes. The proportion of genes with higher *ω* values in the paternal homoeolog than in the maternal homoeolog (*p_PAT_*) minus the proportion of genes with higher *ω* values in the maternal homoeolog than in the paternal homoeolog (*p_MAT_*) is depicted along the x-axis. Mitochondria- (left) and plastid-targeted (right) genes are separated by the degree of interaction: non-interacting genes (top), interacting genes (middle), and genes involved in cytonuclear enzyme complexes (bottom, subset of interacting genes). Proportions are normalized by those found in non-organelle-targeted genes, and genomic bias is denoted by color with maternal bias (i.e., *p_PAT_* – *p_MAT_* < 0) colored purple and paternal bias (i.e., *p_PAT_* – *p_MAT_* > 0) colored green. None of the values exhibited biased proportions according to *χ^2^* tests, relative to genes not targeted to the organelles. Allopolyploids are arranged from oldest (top) to youngest (bottom) as in Figure 2.

**Figure S8.**
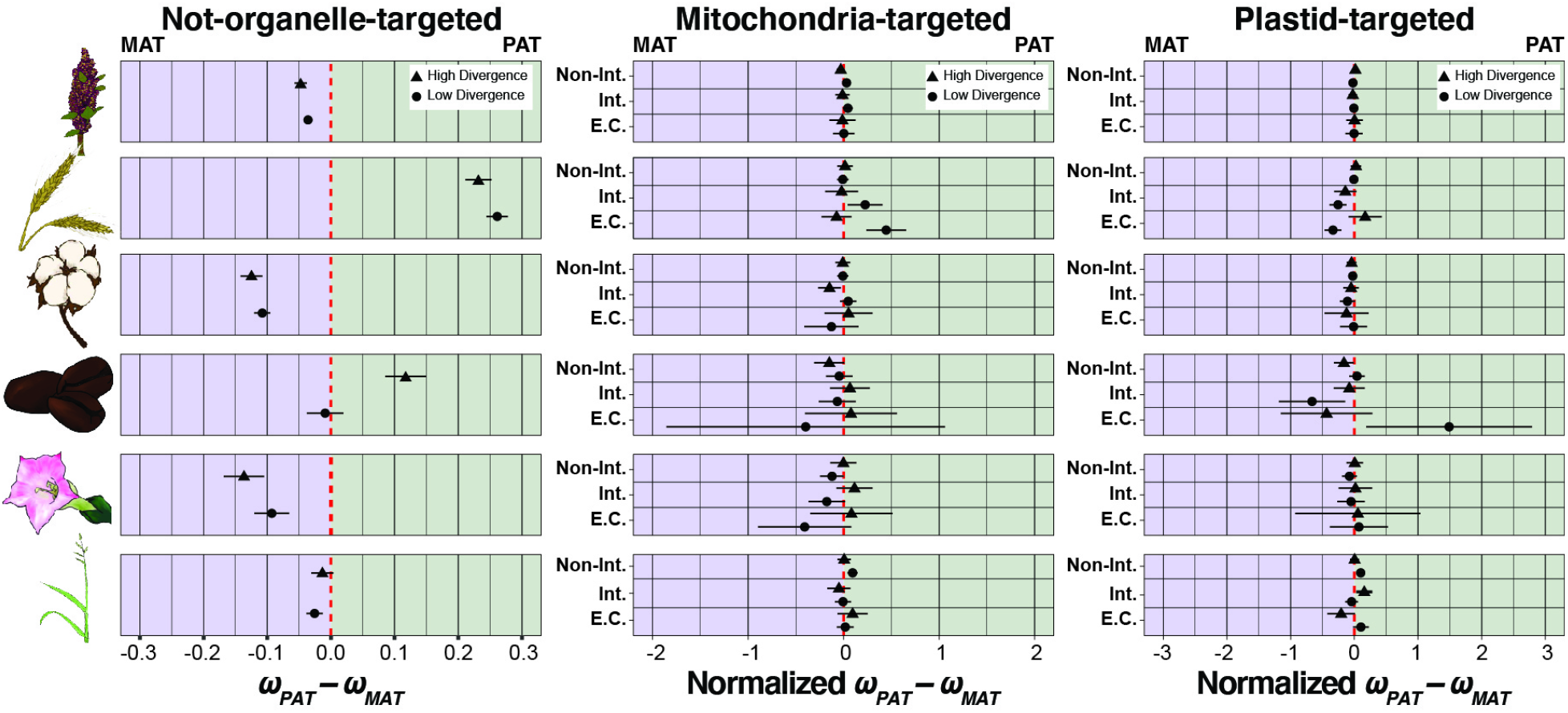
Divergence-binned analysis of subgenomic bias in rates of protein-sequence evolution in six focal allopolyploids. Rates of protein-sequence evolution paternal subgenomes (*ω_PAT_*) minus those found in maternal subgenomes (*ω_MAT_*) are depicted on the x-axis for non-organelle-targeted genes (left), mitochondria-targeted genes (middle), and plastid-targeted genes (right). Genes were concatenated by functional category and binned according to divergence with high-divergence bins (top) depicted by triangles and low-divergence bins (bottom) depicted by circles. Error bars represent standard errors inferred by PAML. Allopolyploids are arranged from oldest (top) to youngest (bottom). The right two panels are further divided by the degree of intimacy of interaction, with non-interacting genes on top, interacting genes in the middle, and genes that are part of enzyme complexes on bottom. The red-dashed line represents equal rates of protein-sequence evolution in the left panel, but on the right two panels the red-dashed line represents the genome-wide pattern taken from the left panel (i.e., organelle-targeted rates were normalized by non-organelle targeted rates). Maternal bias (i.e.,*ω_MAT_* > *ω_PAT_*) occurs left of the red-dashed lines and paternal bias (i.e., *ω_PAT_* > *ω_MAT_*) occurs to the right of the red-dashed line.

**Figure S9.**
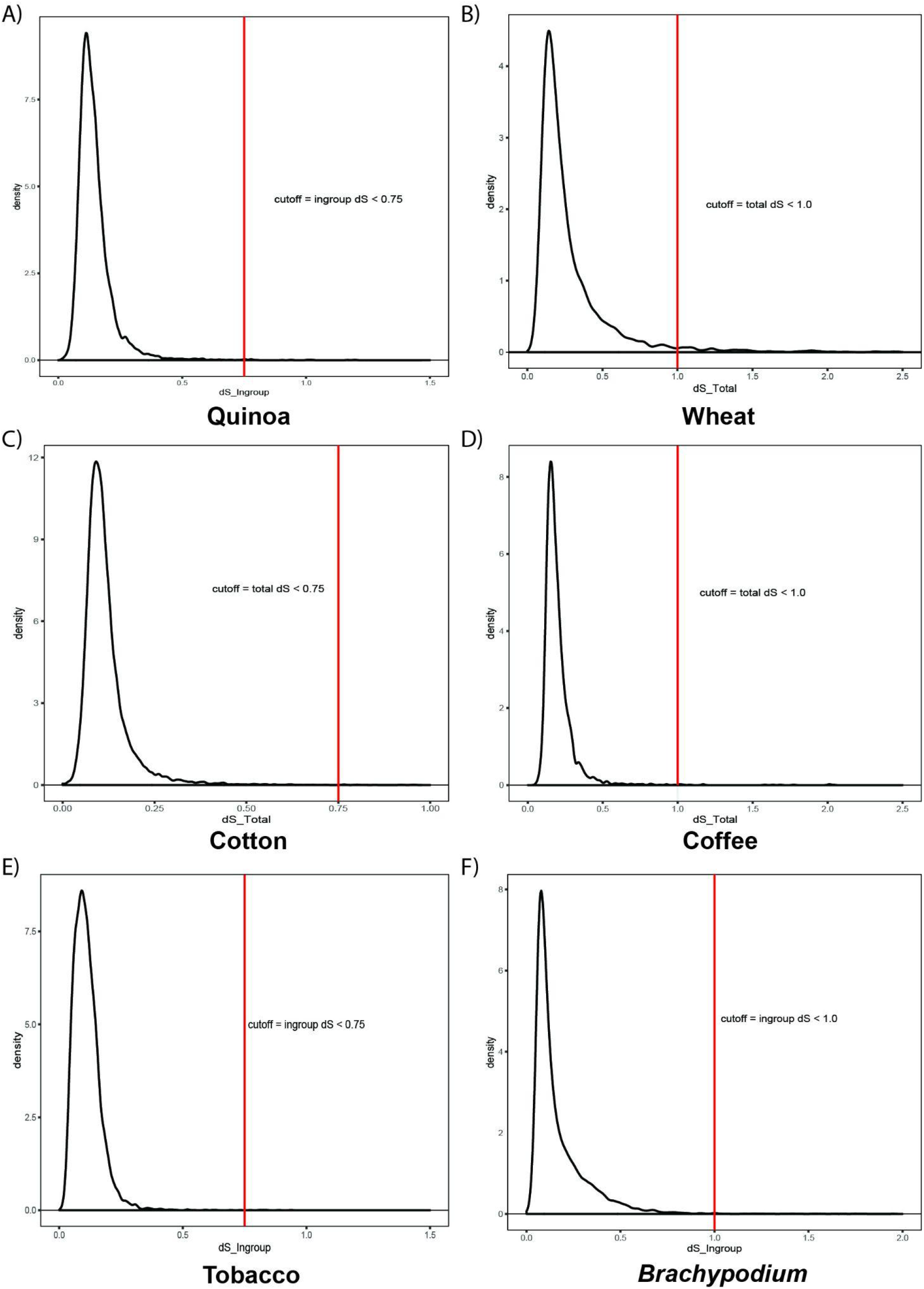
Alignment filtering based on *d_S_* in orthologous quintets. Individual genes with either total *d_S_* values (wheat, cotton, coffee) or ingroup-only *d_S_* levels (quinoa, tobacco, *Brachypodium*) greater than the cutoff point (indicated by the red line) were excluded from analyses of rates of protein-sequence evolution, as we could not exclude the possibility that those quintets were poorly aligned vs. truly divergent.

**Figure S10.**
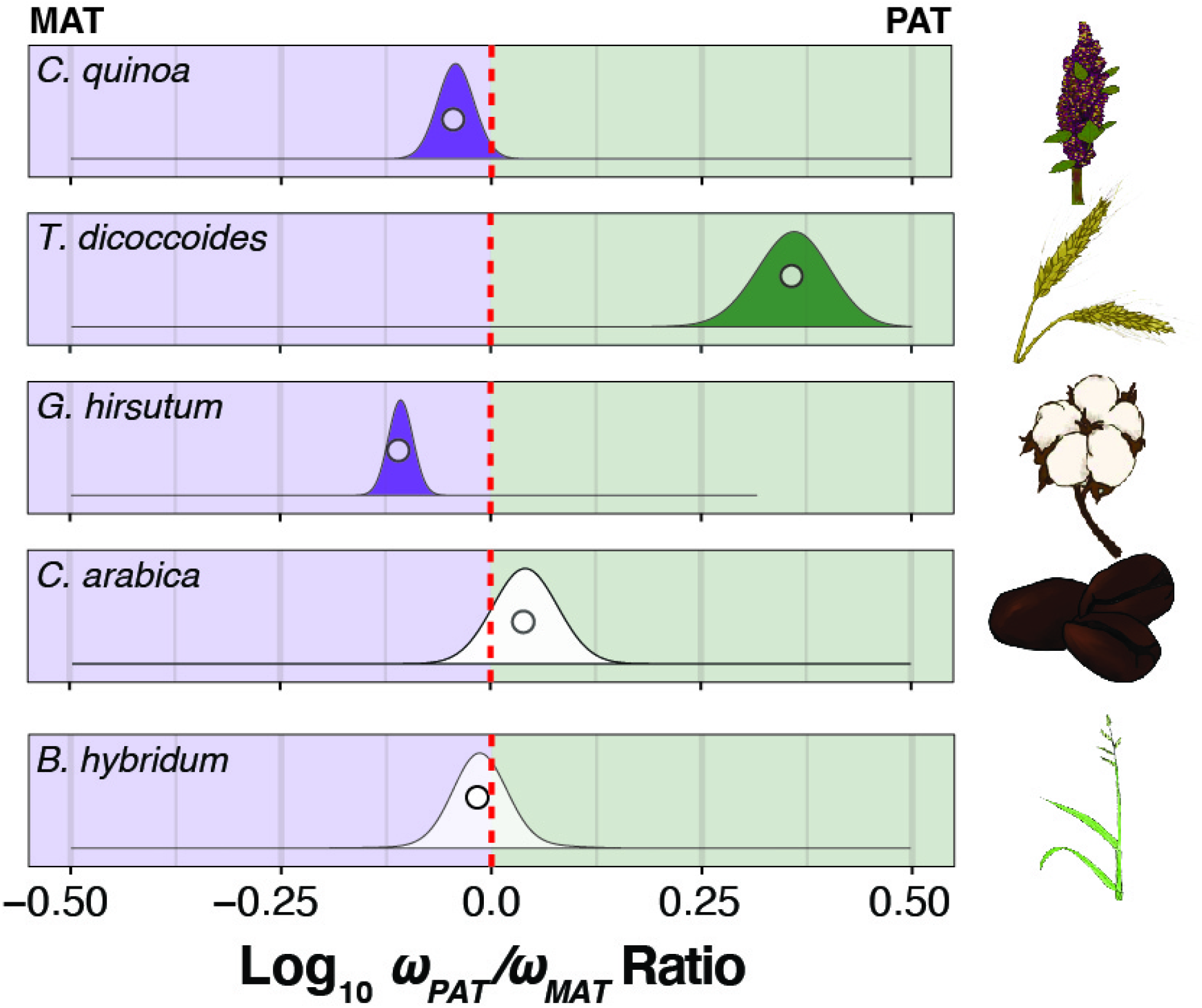
Genome-wide bias in *ω* (*d_N_/d_S_*) across maternal and paternal subgenomes, identical quintets only. Log-transformed ratios of *ω* values in paternal (*ω_PAT_*) vs. maternal (*ω_MAT_*) subgenomes from concatenations (circles), and underlying bootstrap distributions (density curves) of genes encoding proteins that are not targeted to either the plastids or mitochondria using only quintets that were identical across phylogenetic and syntenic methods. Species panels are arranged vertically from oldest (top) to youngest (bottom). Tobacco was excluded from this analysis because it produced so few syntenic quintets. The red-dashed line indicates equal *ω* values across subgenomes, left of the red line indicates higher *ω* values in the maternal subgenomes, and right of the red line indicates higher *ω* values in the paternal subgenome. Bootstrap distributions of *ω* ratios that depart significantly (*p* < 0.05) from the red line are filled in solid according to the direction of subgenomic bias (i.e., green: *ω_PAT_*/*ω_MAT_* > 1.0; purple: *ω_PAT_*/*ω_MAT_* < 1.0; no fill: *ω_PAT_*/*ω_MAT_* ≈ 1.0).

**Figure S11.**
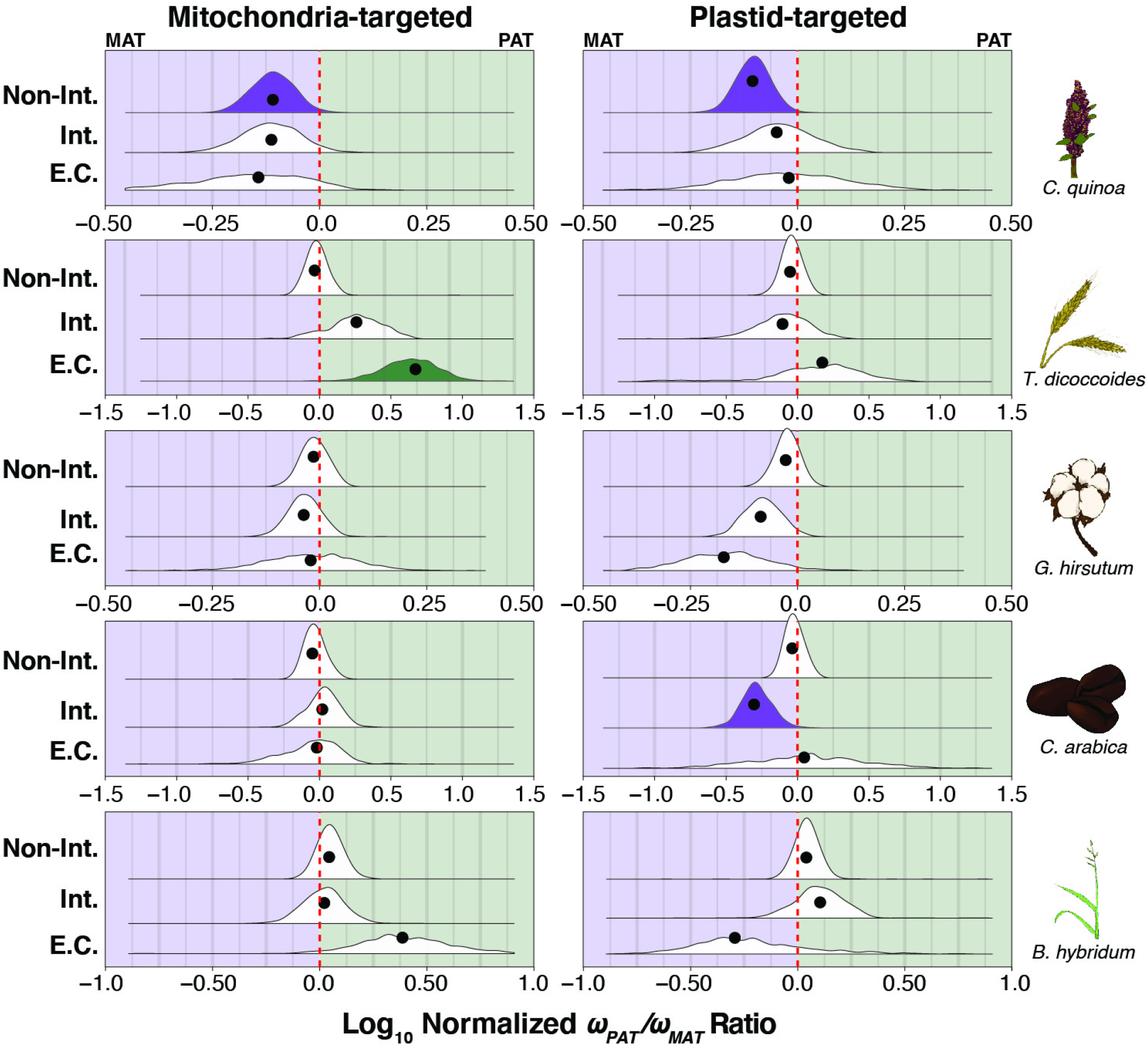
Ratios of maternal vs. paternal *ω* values in organelle-targeted genes, identical quintets only. Log-transformed ratios of maternal vs. paternal *ω* values for concatenations (black circles) and underlying bootstrap distributions (density curves) of mitochondria- (left) and plastid-targeted (right) genes, including only quintets that were identical across phylogenetic and syntenic methods. Species panels are arranged vertically from oldest (top) to youngest (bottom). Tobacco was excluded from this analysis because it produced so few syntenic quintets. The red-dashed line indicates the *ω_PAT_*/*ω_MAT_* ratio for a concatenation of genes not-targeted to the organelles (Figure S9). Ratios left of the red line indicate higher *ω* values in the maternal subgenome, and ratios right of the red line indicate higher *ω* values in the paternal subgenome, after accounting for genome-wide patterns. Bootstrap distributions of *ω* ratios that depart significantly (*p* < 0.05) from the red line are filled in solid according to the direction of subgenomic bias (i.e., green: normalized *ω_PAT_*/*ω_MAT_* > 1.0; purple: normalized *ω_PAT_*/*ω_MAT_* < 1.0; no fill: normalized *ω_PAT_*/*ω_MAT_* ≈ 1.0). The intimacy of interactions are indicated on the y-axis from low or no interaction with organelle gene products (top), to interacting genes (middle), to genes involved in mitochondrial or plastid enzyme complexes (bottom).

## ACKNOWLEDGEMENTS

This work was funded by the National Science Foundations Plant Genome Resources Program (IOS-1829176). We made extensive use of resources from the University of Colorado Boulder Research Computing Group, which is supported by the National Science Foundation (awards ACI-1532235 and ACI-1532236), the University of Colorado Boulder, and Colorado State University. We also thank the Iowa State University ResearchIT Unit for computational support. We thank Jeff Maughan, David Jarvis, Rick Jellen, and Mark Tester for access to the quinoa genomes, discussions of appropriate diploid models, and discussions of orthology inference in the face of homoeologous exchange. We thank Aaron Davis for discussions about polyploidy in *Coffea*, selection of an appropriate outgroup, and a great cup of coffee. We thank John Lovell, Jeremy Schmutz, Pilar Catalan, Sergio Gálvez Rojas, and Robert Hasterok for making the *Brachypodium* genome assemblies available in advance of their recent publication and for general discussions relating to *Brachypodium* genomics. We thank Evan Forsythe and the rest of the Sloan lab for helpful discussion of methods for orthology inference, especially relating to subtree extraction and branch trimming.

