## Supplementary File 1 for "Global patterns of subgenome evolution in organelle-targeted genes of six allotetraploid angiosperms"

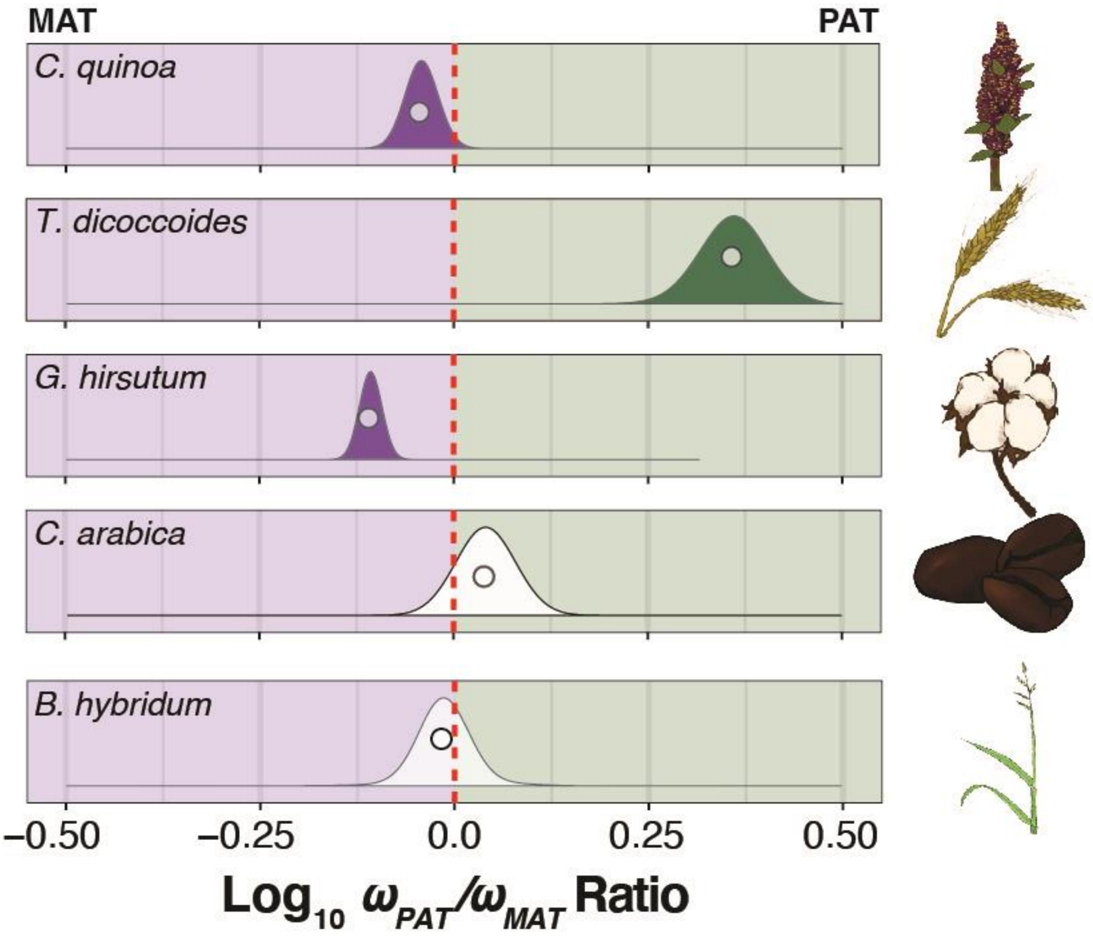


**Figure S10. Genome-wide bias in $\omega$ ($d_N/d_S$) across maternal and paternal subgenomes, identical quintets only.** Log-transformed ratios of $\omega$ values in paternal ($\omega_{PAT}$) vs. maternal ($\omega_{MAT}$) subgenomes from concatenations (circles), and underlying bootstrap distributions (density curves) of genes encoding proteins that are not targeted to either the plastids or mitochondria using only quintets that were identical across phylogenetic and syntenic methods. Species panels are arranged vertically from oldest (top) to youngest (bottom). Tobacco was excluded from this analysis because it produced so few syntenic quintets. The red-dashed line indicates equal $\omega$ values across subgenomes, left of the red line indicates higher $\omega$ values in the maternal subgenomes, and right of the red line indicates higher $\omega$ values in the paternal subgenome. Bootstrap distributions of $\omega$ ratios that depart significantly ($p < 0.05$) from the red line are filled in solid according to the direction of subgenomic bias (i.e., green: $\omega_{PAT}/\omega_{MAT} > 1.0$; purple: $\omega_{PAT}/\omega_{MAT} < 1.0$; no fill: $\omega_{PAT}/\omega_{MAT} \approx 1.0$).

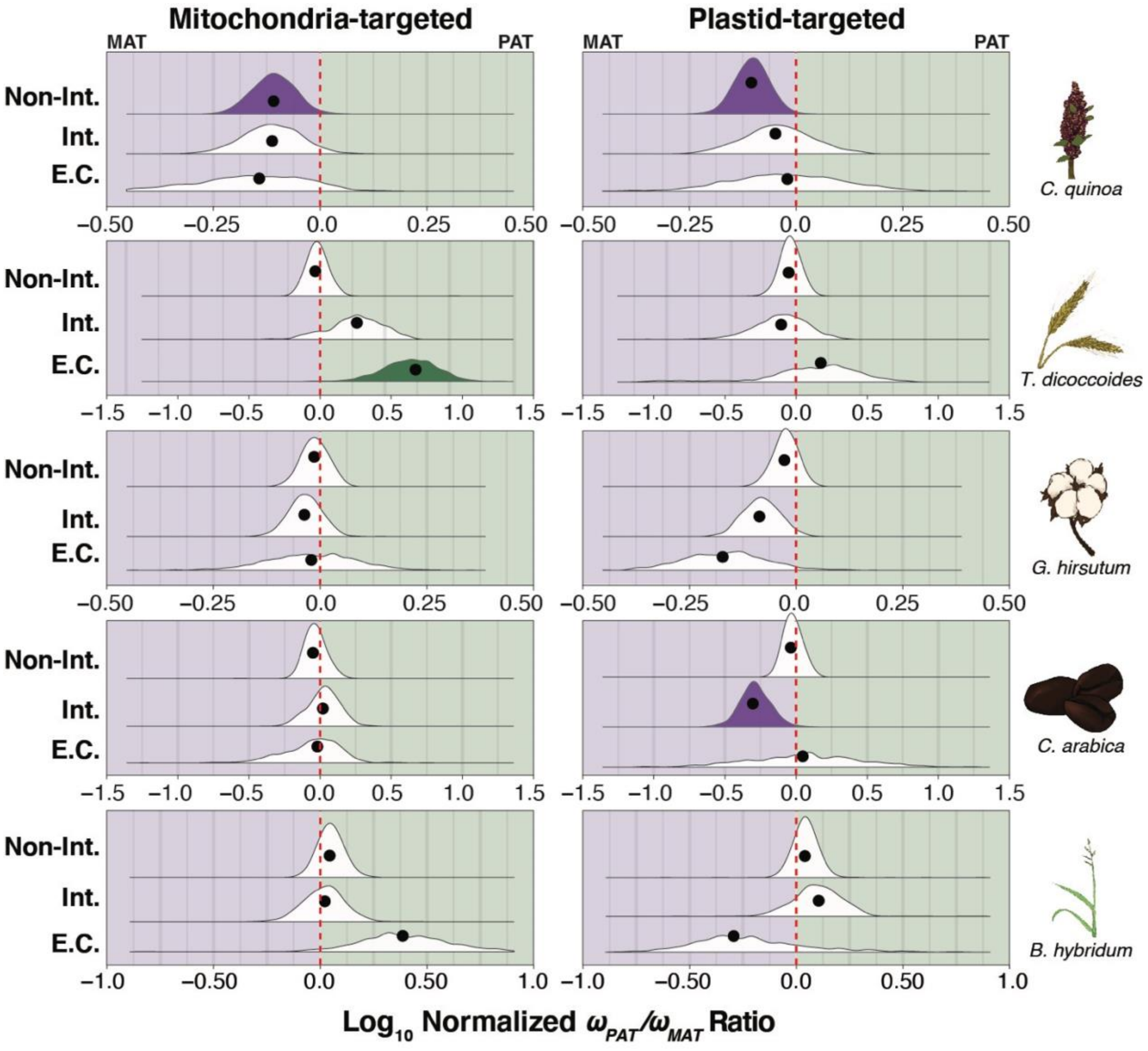


**Figure S11. Ratios of maternal vs. paternal $\omega$ values in organelle-targeted genes, identical quintets only.** Log-transformed ratios of maternal vs. paternal $\omega$ values for concatenations (black circles) and underlying bootstrap distributions (density curves) of mitochondria- (left) and plastid-targeted (right) genes, including only quintets that were identical across phylogenetic and syntenic methods. Species panels are arranged vertically from oldest (top) to youngest (bottom). Tobacco was excluded from this analysis because it produced so few syntenic quintets. The red-dashed line indicates the $\omega_{PAT}/\omega_{MAT}$ ratio for a concatenation of genes not-targeted to the organelles (Figure S9). Ratios left of the red line indicate higher $\omega$ values in the maternal subgenome, and ratios right of the red line indicate higher $\omega$ values in the paternal subgenome, after accounting for genome-wide patterns. Bootstrap distributions of $\omega$ ratios that depart significantly ($p < 0.05$) from the red line are filled in solid according to the direction of subgenomic bias (i.e., green: normalized $\omega_{PAT}/\omega_{MAT} > 1.0$; purple: normalized $\omega_{PAT}/\omega_{MAT} < 1.0$; no fill: normalized $\omega_{PAT}/\omega_{MAT} \approx 1.0$). The intimacy of interactions are indicated on the y-axis from low or no interaction with organelle gene products (top), to interacting genes (middle), to genes involved in mitochondrial or plastid enzyme complexes (bottom).
